# Rapid spectrophotometric detection for optimized production of landomycins and characterization of their therapeutic potential

**DOI:** 10.1101/2023.11.07.566088

**Authors:** Todd C. Chappell, Kathleen G. Maiello, Allison J. Tierney, Karin Yanagi, Jessica A. Lee, Kyongbum Lee, Charles R. Mace, Clay S. Bennett, Nikhil U. Nair

**Affiliations:** Department of Chemical & Biological Engineering, Tufts University, Medford, MA; Department of Chemistry, Tufts University, Medford, MA

**Keywords:** Keywords, Polyketide, angucycline, media optimization, *Streptomyces*, natural product, chemotherapeutic

## Abstract

Microbial derived natural products remain a major source of structurally diverse bioactive compounds and chemical scaffolds that have potential as new therapeutics to target drug resistant pathogens and cancers. In particular, genome mining has revealed the vast number of cryptic or low yield biosynthetic gene clusters in the genus *Streptomyces*. Here, we describe our efforts to improve yields of landomycins – angucycline family polyketides under investigation as cancer therapeutics – by a genetically modified *Streptomyces cyanogenus* 136. After simplifying the extraction process from *S. cyanogenus* cultures, we identified a wavelength at which the major landomycin products absorb in culture extracts, which we used to systematically explore culture medium compositions to improve total landomycin titers. Through correlational analysis, we simplified the culture optimization process by identifying an alternative wavelength at which culture supernatants absorb yet is representative of total landomycin titers. Using the subsequently improved sample throughput, we explored landomycin production during the culturing process to further increase landomycin yield and reduce culture time. Testing the antimicrobial activity of the isolated landomycins, we report broad inhibition of Gram-positive bacteria, inhibition of fungi by landomycinone, and broad landomycin resistance by Gram-negative bacteria that is likely mediated by exclusion of landomycins by the bacterial membrane. Finally, the anticancer activity of the isolated landomycins against A549 lung carcinoma cells agrees with previous reports on other cell lines that glycan chain length correlates with activity. Given the prevalence of natural products produced by *Streptomyces*, as well as the light-absorbing moieties common to bioactive natural products and their metabolic precursors, our method is relevant to improving the yields of other natural products of interest.

## Introduction

Advances in higher throughput and more sensitive technologies for the identification of natural products and biosynthetic gene clusters (BGCs) – genes that encode enzymatic pathways responsible for natural product biosynthesis – have revealed a vast landscape of natural biochemical diversity.^1,2^ Natural products, especially those with complex polycyclic structures, continue to serve as promising new drug leads to treat several relevant health threats, such as multidrug resistant cancers and antibiotic resistant microbial pathogens.^3–5^ Though biosynthesis is a catalytically efficient method to produce natural products, BGC expression is often poor and pathway activation mechanisms remain indeterminant. Additionally, while efforts to develop chemical synthesis methods have met with significant success, they are often expensive, time-consuming, and low-yielding, and thus are better suited for semi-synthetic methods that start with biologically derived chemical scaffolds.^6^ Thus, production of sufficient bioactive natural products remains a major hurdle that limits their eventual translation to clinical applications.

Of the thousands of identified microbial BGCs, the genus *Streptomyces* encodes the greatest number and highest density, producing numerous bioactive compounds and relevant chemical scaffolds.^7–9^ Among these, landomycins – polyketides that belong to the angucycline family – exhibit bioactivity via diverse and potentially novel mechanism(s) of action, making them promising therapeutics to target drug-resistant phenotypes.^10–13^ Landomycin BGCs have only been directly identified in three genomic samples, though a fourth is presumed due to landomycin detection in culture extracts.^14–17^ Landomycin A (lanA) and landomycin E (lanE) are the most widely-studied landomycins, and both possess the widest reported spectrum of anticancer activities among the landomycins.^18^ The structures of lanA and lanE only differ in the composition of their glycan chain. LanA has the longest glycan of naturally occurring landomycins, a hexasaccharide composed of a repeated trisaccharide unit of two D-olivoses and a single L-rhodinose, while lanE, contains only a single trisaccharide unit.^10,18,19^ Very limited data is available regarding the antimicrobial spectra of landomycins, with only a few studies reporting activity against one or two sensitive organisms.

*Streptomyces cyanogenus* 136 (Sc136) is the only known natural source of lanA.^20^ Though meticulous and innovative methods have demonstrated the feasibility to chemically synthesize the complex benz[*a*]anthracene core, hexasaccharide glycan, complete lanA molecule, and a variety of non-natural landomycins, these methods are not suitable for the production of landomycins on the scale required for drug development.^21–30^ Alternatively, control over the biosynthetic regulon via genetic interventions, including genetic modifications to Sc136, cloning the lanA BGC into another *Streptomyces*, and optimization of culture conditions, have met with substantial success at improving yield or altering dominant products.^14,31–34^ In particular, modifications to the major pathway regulators *adpA* and *bldA* have improved landomycin titers – though modifications to pathway gene expression can alter the dominant and minor landomycin products compared to wildtype Sc136.^35–37^ Of particular note is recent work that showed that complementation of Sc136 with a functional copy of the transcription factor *adpA* from *S. ghanaensis* (*adpA_gh_*) improved lanA yields by more than 5-fold and rendered Sc136 capable of synthesizing lanA in a variety of media.^36^ Still, additional improvements to increase the yield, as well as methods that facilitate the purification of landomycins, will aid in their potential for future clinical testing and modification into next-generation, semi-synthetic analogs unavailable via biosynthesis.

Here we describe the development and utilization of spectrophotometric methods to optimize landomycin production, and the characterization of the antimicrobial and anticancer activity of the major landomycin products of an *adpA_gh_* expressing Sc136 strain (Sc92a). To this end, we characterized the major landomycin products of Sc92a, improved extraction and fractionation conditions, and identified a wavelength of light (265 nm; A265) at which all of these products absorb. Using A265 as an estimate for total landomycin production, we systematically explored the impact of media components on landomycin titers. We further simplified and increased the throughput of this method by correlating the absorbance measurements from culture extracts to those of supernatants, ultimately identifying a wavelength (345 nm) where the culture supernatant absorbance is representative of landomycin titers. Of the microbes tested, we found that Gram-positive bacteria are most sensitive to landomycins and fungi are inhibited by landomycinone, although mostly at concentrations that are unlikely to be clinically relevant. Gram-negative organisms are resistant to landomycins but could be sensitized by disrupting the outer membrane. Consistent with prior studies, we confirm that the landomycins exhibit potent anticancer activity against lung carcinoma cells (A549), indicating the future potential to develop landomycins into effective anti-cancer chemotherapeutic candidates.

## Materials and Methods

### Culture conditions

Strains used in this study are listed in **Table S1**. *S. cyanogenus* was grown in TSB for general propagation. *Bacillus subtilis*, *Listeria seeligeri*, *Lactococcus cremoris*, *Enterococcus faecium*, *Staphylococcus aureus*, *E. coli*, *Pseudomonas aeruginosa*, and *Salmonella enterica* ser. Typhimurium were cultured in Brain Heart Infusion (BHI) broth. *Saccharomyces cerevisiae*, and *Candida albicans* were cultured in 2×YPD (40 g/L casein peptone, 20 g/L yeast extract, 20 g/L glucose, 100 mg/L adenine). For solid media growth, 15 g/L agar was added to BHI or 2×YPD. All cultures were grown at 250 rpm shaking and 37 °C, except for *S. cyanogenus*, *L. cremoris*, *S. cerevisiae*, and *C. albicans*, which were grown at 30 °C.

### Landomycin production and mycelia fractionation

Initial *S. cyanogenus* propagation and landomycin production conditions were kindly provided by personal communication from Prof. Bohdan Ostash (Ivan Franko Lviv National University, Lviv, Ukraine). *S. cyanogenus* 136 +pOOB92a (a.k.a. Sc92a) frozen stock was streaked on soy mannitol agar (SMA; 20 g/L soy flour, 20 g/L mannitol, 15 g/L agar, pH to 8.0) and grown for 48 h. Colonies with a red-brown halo were used for preculture. A 1–2 cm^2^ piece of agar with visibly grown colonies was cut from the plate and added to 30 mL of SG medium (20 g/L glucose, 10 g/L phytone peptone, 2 g/L CaCO_3_, pH to 7.0; autoclave; add 1.9 mg/L CoCl_2_·6H_2_O) in a 250 mL flask and grown for 48 h as preculture. 2.5 mL of preculture was used to inoculate each of 10 × 500 mL flask with 100 mL SG and grown for 48 h as the production culture. SG was modified as required for medium optimization by adding or replacing components prior to autoclaving (nitrogen sources), or adding concentrated stocks after autoclaving (glucose, metal ions). The culture solids were pelleted by centrifugation (4500 ×g for 10 min) and the supernatant decanted, buffered to pH 7.0, and set aside for extraction. The cell-mycelium and culture solids pellet were then resuspended in PBS, vortexed vigorously to homogenize, and centrifuged (4500 ×g for 5 min) again. The supernatant was decanted and set aside for extraction. The pellets were again resuspended to 35 mL total volume in PBS and vortexed vigorously. The homogenized samples were then centrifuged for 1–1.5 min at 4500 ×g to selectively pellet the cell-mycelium fraction, while retaining the culture solids in suspension to decant for extraction. Resuspension, vortexing, and centrifugation steps were repeated until the pellet was composed only of a beige-grey cell-mycelium pellet, and the supernatant was clear.

### Isolation of pure landomycins

Initial isolation methods were determined from previous works and adapted as needed.^10,19^ The supernatant fraction (∼1 L) from the *S. cyanogenus* 136 +pOOB92a was extracted once with 1 L and twice with 500 mL ethyl acetate. The 500 mL culture solids suspension was extracted with three 1 L, followed by two 500 mL volumes of ethyl acetate. The combined extract was evaporated down to solid crude material and put under high vacuum to ensure dryness.

The crude red solid (904.2 mg) was dissolved in about 5 mL of 7% MeOH/CHCl_3_ solution and loaded on to a Yamazen column (180 g of silica) equilibrated for two minutes at 3% MeOH/CHCl_3_. The system used was an Automated flash column chromatography (normal phase) Smart Flash EPCLC W-Prep 2XY Dual Channel Automated Flash Chromatography System with an additional ELSD detector, provided by Yamazen Corporation. The column was run at 4% MeOH/CHCl_3_ for twenty minutes then increased over two minutes to 10% MeOH/CHCl_3_ to run for an additional fifteen minutes. This column yielded fractions I (pink/purple solid), II (red solid), and III (orange/red solid). Fraction I was evaporated to dryness, then separated further by size exclusion chromatography with 40 g LH-20 Sephadex (Sigma Aldrich), in 50% MeOH/CHCl_3_ (2 × 50 cm) to yield pure landomycinone (6.7 mg) and 5,6-anhydrolandomycinone (0.3 mg). Fraction II was evaporated to dryness, then precipitated from a concentrated chloroform solution into pentane, giving pure landomycin A (423.1 mg). Fraction III was evaporated to dryness, then separated further with 40 g LH-20 Sephadex in MeOH (2 × 50 cm) to yield landomycin B and landomycin D. After rotary evaporation, landomycin B (39.2 mg) was precipitated from a concentrated chloroform solution into pentane. Landomycin D (9.6 mg) was precipitated from a concentrated ethyl acetate solution into pentane. Products were visualized on TLC using UV and by staining with a 5% aqueous sulfuric acid solution. The purity of all landomycin species was confirmed by a normal phase Elite LaChrom Hitachi HPLC (90% EtOAc in Hexanes). NMR spectra were recorded on a Bruker Avance III NMR spectrometer at 500 MHz for ^1^H NMR and 125 MHz for ^13^C-NMR. Mass spectrometry data was collected on a low resolution Finnigan LTQ ESI-MS (ThermoFisher). See Supplementary Information for more details. All solvents used were HPLC grade (ThermoFisher Scientific).

### Landomycin standard curves

All standard curves were generated by diluting landomycin stocks (10 mg/mL in DMSO) 10× into Dulbecco’s PBS (DPBS, pH 7.0; Corning), followed by 3-fold serial dilutions into DPBS 10% DMSO, and performing an absorbance spectral scan (200–1000 nm; 5 nm increments) using a Spectramax M3 (Molecular Devices).

### Spectrophotometric detection of landomycins from culture extracts and supernatants

Absorbance of culture supernatants was determined by pelleting culture solids from the landomycin production cultures and diluting the resulting supernatant 2× into DPBS or 10× into HEPES buffer (100 mM, pH 7.0). Extraction of landomycin fractions for spectrophotometric determination of relative concentrations was performed by adding 300–500 µL aliquots of landomycin culture (supernatant, cells, and mycelia) to an equal volume of chloroform. Samples were then vortexed for 30 s and centrifuged to separate fractions. Samples were vortexed and centrifuged for a second time and a 100 µL aliquot of the chloroform extract was removed from the lower fraction. Samples were dried at room temperature under vacuum for approximately 30 min using an Eppendorf Vacufuge and stored at –80 °C. Immediately before recording absorbance readings, samples were resuspended in 100 µL DMSO and diluted 10× into DPBS. Spectrophotometric scans from 200–800 nm at 5 nm intervals were performed for all supernatant and extract samples.

### General A549 cell culture

Human non-small cell lung cancer (NSCLC) cells (A549; CCL-185) from ATCC were cultured in F-12K medium (ATCC) with 10 % (v/v) fetal bovine serum (Neuromics). Cultures of A549 cells were incubated at 37 °C, 95 % humidity, and 5 % CO_2_ atmosphere. Cells were passaged every three to four population doubling times, when their concentration was approximately 4 × 10^6^ cells mL^−1^ (approximately 3 days between each passage). All experiments were performed using cultures that had been passaged less than twenty times.

### A549 viability assay

A549 cells were seeded at 10,000 cells per well (200 µL) in 96-well tissue culture treated plates (CellTreat) (t = 0 h). 20 mM concentrated stocks of landomycins and doxorubicin hydrochloride (Tocris Biosciences) were made in DMSO immediately prior to addition to cells. Concentrated stocks to 20 mM were diluted to make working stocks that were diluted to 0–100 µM in cell culture medium with a final DMSO concentration of 0.5 % (v/v). At 24 h, the medium was aspirated from each well, washed with 100 µL PBS, and 0–100 µM of product of interest was added to each well in complete media (supplemented with a final concentration 0.5% v/v DMSO), and incubated cells for 24 h. This concentration of DMSO afforded no change in viability when assessed without drug present and compared to the 0 µM sample (> 95% cell viability with 0.05% DMSO). Then, the medium was aspirated, washed cells with PBS, and trypsinized cells using 100 µL of 0.25% trypsin/EDTA (Gibco) to release the cells from the surface of the plate. Trypsin/EDTA was neutralized with an equal amount (100 µL) of complete media. Cells were pelleted (300 × g, 8 minutes), and washed with 100 µL PBS. Three stains were used to classify the viability of cells: Hoechst 33342 to identify nuclei, and Annexin V-FITC, and Ethidium homodimer I to label viable, apoptotic, and necrotic cells, respectively. 100 µM concentrated stocks of Ethidium homodimer I and Hoechst 33342 were prepared in PBS. Then, 5 µM working solutions of Annexin V-FITC (BioLegend), Ethidium homodimer I (Biotium), and Hoechst 33342 (Thermo Scientific) were prepared in 1× Annexin V Binding Buffer (100 mM HEPES, 25 mM CaCl_2_, 1.4 M NaCl; BioLegend). To analyze cells, 50 µL of Annexin V-FITC, Ethidium homodimer I, and Hoechst 33342 working solutions was added to each well containing a washed cell pellet. An additional three wells received each of the single dyes for compensation controls. Samples were incubated in the well plate for 10 minutes (37 °C, 5% CO_2_). Prior to plate analysis via flow cytometry, 200 µL of 1× Binding Buffer was added to each well. All wells were analyzed using a Guava easyCyte 12HT flow cytometer (Cytek). Different laser/filter combinations were used for each dye (**Table S2**). Gain controls were adjusted for GRN-B, RED-B, and BLU-V channels to 1.00 to accommodate dyes used (**Table S3**), and YEL-B (emission filter 575/25) was also adjusted to 1.00 to accommodate the natural fluorescence of doxorubicin hydrochloride (ex/em: 470/560).

### Agar well diffusion assay

Landomycin stock solutions were made by resuspending purified landomycin solid fractions at 10 mg/mL (lanD, lanB, lanA, lan, and A-lan) in DMSO. Stock solutions were stored at –20 °C. Overnight cultures of each strain were diluted 50× into fresh medium and 100 µL was spread-plated on the appropriate solid agar medium. Agar plates were allowed to dry and circular wells were excised. Working solutions of landomycin were made by diluting concentrated stocks to 250 ppm (µg/mL) into PBS pH 7.0, and 100 µL of a working solution was added to an agar well. Plates were imaged after overnight incubation at the appropriate temperature.

### Microbial MIC assay

Overnight cultures of indicated strains were sub-cultured 1:1000 (bacteria) or 1:500 (fungi) into fresh media containing 250 µg/mL landomycin (diluted from 10 mg/mL stocks), or 2.5% (v/v) DMSO (control) and aliquoted into 96-well plates. Subcultures were serially diluted 2-fold into fresh medium with 2.5% DMSO to ensure consistent vehicle concentration. Plates were sealed with parafilm to prevent culture evaporation and grown in a Biotek Epoch 2 microplate reader (37 °C, continuous linear shaking at 731 cpm) for 24–48 h, with an absorbance reading at 600 nm taken every 5 min. Due to the change in absorbance over time caused by the breakdown of the landomycins, cultures were blanked at each time point using BHI media with an equivalent concentration of each landomycin.

### Statistics

Standard linear regression was performed using GraphPad Prism and Microsoft Excel. Ordinary one-way ANOVA and correlational analysis (Pearson’s r) were performed using GraphPad Prism.

**Figure 1:**
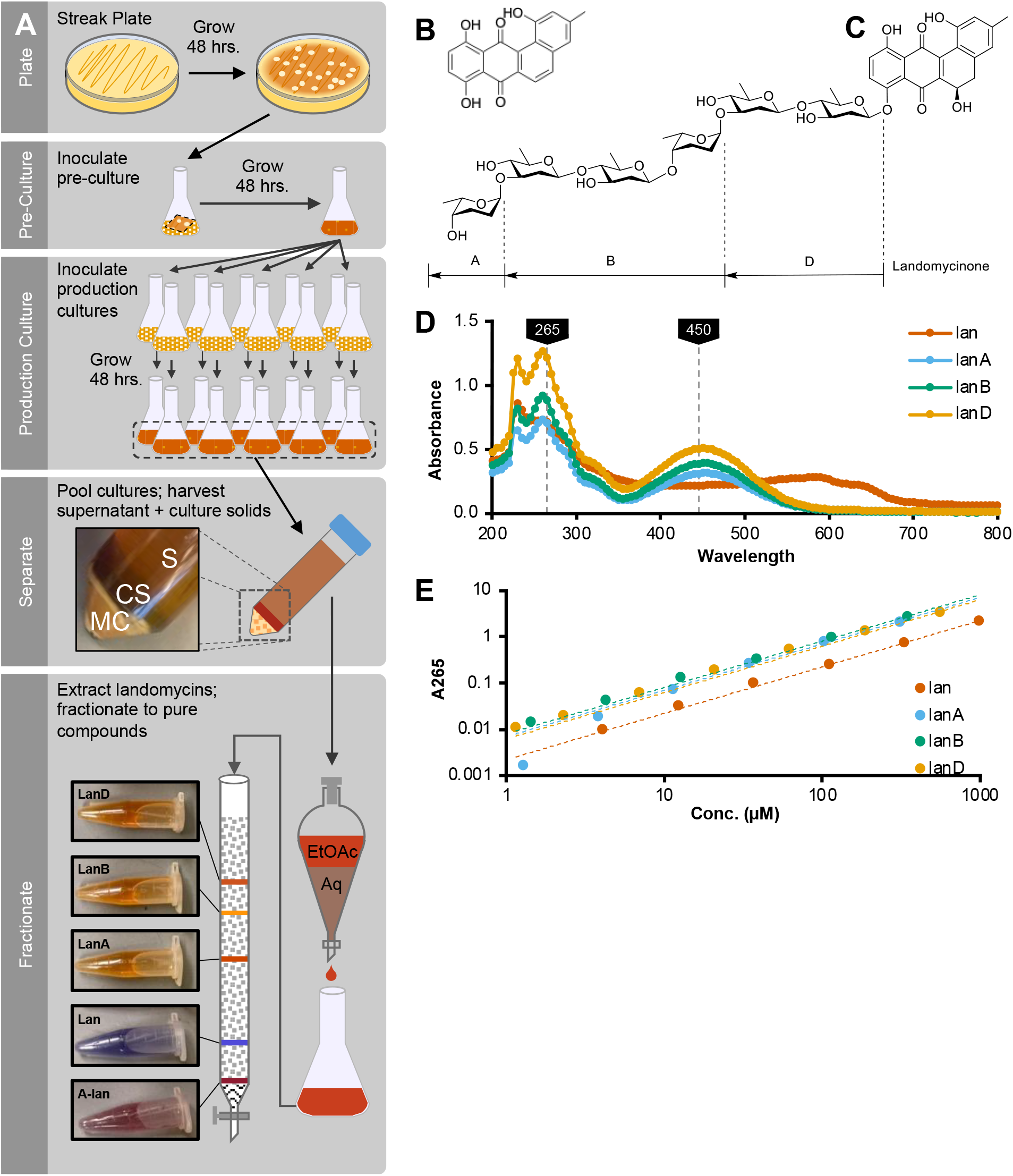
Landomycin production method and major products of *S. cyanogenus* 136 +pOOB92a. A) The major steps of landomycin production include i) differentiating Sc136 on SMA agar, ii) inoculating a preculture using an agar plug, and iii) subculturing the preculture into production cultures. Culture supernatants (S) and (CS) solids are then separated from cells and mycelium (MC) and landomycins are extracted from the aqueous using ethyl acetate (EtOAc), and fractionated on a silica column. B&C) Structures of 5 major products, including: i) anhydrolandomycinone (A-lan), ii) landomycinone (lan), iii) landomycin A (lanA), iv) landomycin B (lanB), and v) landomycin D (lanD). D) Absorbance spectra of purified lan, landomycin A (lanA), landomycin B (lanB), and landomycin D (lanD). Absorbance peaks at 265 and 450 nm are labeled. Readings were taken every 5 nm from 200-800 nm. E) Standard curves and absorbance spectra for lan, lanA, lanB, and lanD demonstrating linear relationship of absorbance to compound concentration (R^2^ ≥ 0.99 for all trendlines).

## Results

### An improved extraction and isolation protocol for landomycin products

Landomycin production by *Streptomyces* is commonly performed in a three-step process: i) differentiation on agar plates, ii) preculturing, and iii) production culturing (**Fig. 1A**).^19^ Landomycins are then extracted from production cultures using a biphasic extraction, and the extract is fractionated using traditional separation methods to isolate individual components. Though landomycins can be extracted quite simply from the supernatant, this approach sacrifices much of the landomycin produced (>80%), which remains trapped in culture solids. Previous reports of extraction from total cultures (cells+mycelia+solids+supernatant) have been reported to produce higher yields, however, our initial efforts produced greasy, poorly crystalized solids that retained a considerable amount of contaminant.^19,34^ To improve product quality and reduce the complexity of separations, we developed a simple method to fractionate the colored culture solids, likely densely packed with secondary metabolites such as landomycins, from the remaining cells and/or mycelia. We predicted that the size and density difference between cells/mycelia and colored culture solids would be sufficient to separate by differential centrifugation, and we found that with successive rounds of vortexing and pelleting generated two distinct layers of well separated solids: i) a more dense, white-tan layer (cells+mycelium) on the bottom and ii) a less dense, red-brown layer (culture solids) on the top (**Fig. 1A, Fig. S1**). Using the extracts from the supernatant and culture solids fractions, we were able to simplify the previously reported fractionation process to a single normal phase silica column and a precipitation step to obtain still obtain 390–430 mg/L lanA at high purity (>97%). We identified 5 major products in Sc92a cultures: anhydrolandomycinone (A-lan), landomycinone (lan), landomycin A (lanA), landomycin B (lanB), and landomycin D (lanD) (**Fig. 1A–C**) .

### Optimized media formulations enhance landomycin production

The electron orbitals of polyketides are frequently oriented such that electrons can absorb light in the near UV (>200 nm) to near infra-red (< 800 nm) range, which is detectable by standard UV-vis spectroscopy. More specifically, lanA production by Sc136 has been estimated by measuring the absorbance of culture extracts at 445 nm.^36^ We therefore explored the applicability of a similar spectrophotometric approach to detect the other soluble landomycin products that we identified in culture extracts (lan, lanA, lanB and lanD). To validate this approach, we acquired absorbance spectra of the purified compounds from 200–800 nm (**Fig. 1D**). We found a wide absorbance peak from approximately 350–550 nm, with a maximum at 450 nm, which was specific to the glycosylated forms of landomycin (lanA, lanB, and lanD). Conversely, lan absorbed well from 575–650 nm, while the glycosylated forms had negligible absorbance in this range. All compounds absorbed well in the UV range, with peak maxima at 230 and 260 nm, suggesting absorbances near these wavelengths could be effective determinants of total lan, lanA, B, and D production. We generated standard curves using linear regression for lan, lanA, lanB, and lanD at 265 nm and found a strong goodness of fit for all compounds (R^2^ ≥ 0.99) (A265; **Fig. 1E**).

Integration of *adpA* in Sc136 alters the conditional expression of the landomycin biosynthetic pathway in diverse media. This alleviates the dependence on the standard soytone-glucose (SG) production medium for landomycin production and presents an opportunity to improve production by optimizing the medium composition. SG is composed of a soybean derived peptone (soytone or phytone peptone), glucose, calcium carbonate and cobalt, though the causal components of SG essential to landomycin production are unknown.^19^ Accordingly, we utilized our newly identified correlation between A265 and the concentration of the major landomycin products to approximate total landomycin production in culture extracts, thereby identifying production limitations and improving landomycins yield from Sc92a cultures in modified SG compositions.

**Figure 2:**
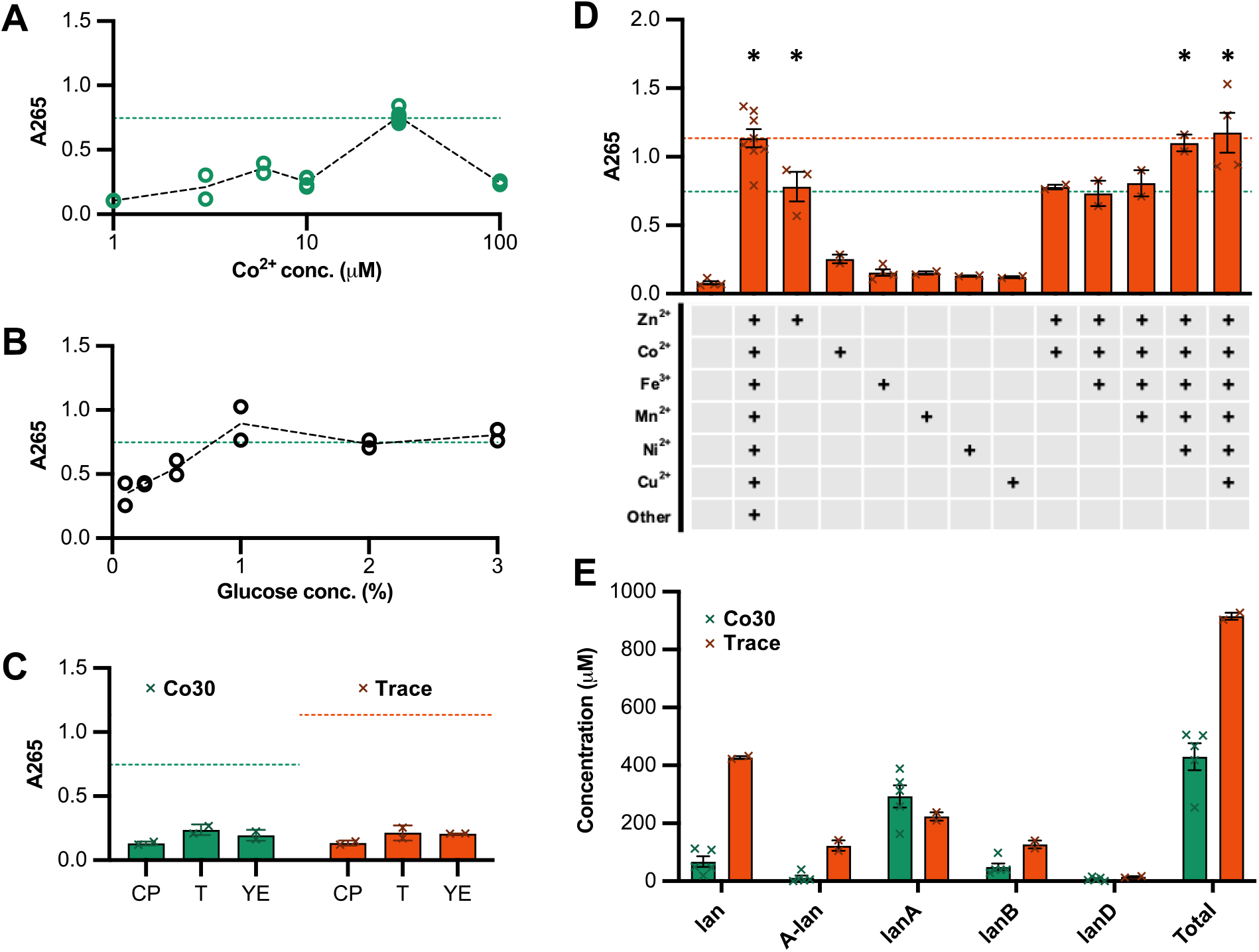
Optimization of landomycin production medium. Chloroform extractions of cultures with modified media compositions were tested to determine the impact on landomycin production A) Effect of cobalt concentration on landomycin production in SG medium. Concentrations tested include 1, 3, 6, 10, 30 and 100 μΜ. B) Effect of glucose concentration on landomycin production in SG medium with 30 μΜ Co. Concentrations tested include 0.1, 0.25, 0.5, 1, 2, and 3% (w/v) glucose C) Effect of replacing phytone peptone with casein peptone (CP), tryptone (T), or yeast extract (YE) in the presence of 30 μΜ Co (green) or trace metals solution (orange). D) Impact of trace metal composition in SG medium on landomycin production. Metal ions were added as the following salts from 1000x stocks (final concentrations given): 10 μM ZnCl_2_, 2 μM CoCl_2_, 50 μM FeCl_3_, 10 μM MnCl_2_, 2 μM NiCl_2_, 2 μM CuSO_4_. Other indicates the use of a commercial trace metals solution which additionally contains 2 μM Na_2_MoO_4_, 2 μM Na_2_SeO_3_, 20 μM CaCl_2_, and 2 μM B(OH)_3_. * denotes samples with means significantly different from all other unlabeled means and not significantly different from one another, as determined by ordinary one-way ANOVA and Tukey multiple comparison test (p<0.05). E) Landomycin yield from 1 L production cultures of Co30 (green) or Trace (orange) media. Error bars represent SEM.

We first investigated the effect of cobalt concentration on landomycin production (**Fig. 2A**). SG medium contains 0.001 g/L (7.7 µM) cobalt, yet we found that increasing cobalt to 30 µM increased total landomycins by ∼2-fold. Increasing cobalt concentration further to 100 µM resulted in a drop in landomycin production, and cultures turned a dark brown-black, which is frequently associated with poor yields that are thought to be due to degradation of landomycins. Using the improved 30 µM cobalt concentration in the SG base medium (Co30), we next investigated the effect of glucose concentration on landomycin production (**Fig. 2B**). Relatively high glucose concentrations (2 %) are cited as necessary for good landomycin production, potentially due its role as a source of reducing potential and as a metabolic precursor to glucose-6-phosphate, both of which are required to generate the constituent deoxygenated sugars, olivose and rhodinose. However, glucose is also known to play an important role regulating secondary metabolism, frequently repressing pathways while nutrients are abundant. We found that glucose is required at a minimum concentration of 1.0 % (w/v) to obtain maximum landomycin titers and there was no detriment to further increasing its concentration, indicating that production is not glucose inhibited or limited (**Fig. 2C**). Maintaining 2 % glucose, we then investigated different complex nitrogen sources but found that they all resulted in a drastic reduction in landomycin titer. This result suggests that specific components of the phytone peptone are necessary to achieve maximum landomycin yields. Finally, since previous studies have had success in replacing cobalt with other divalent metal ions, we tested the suitability of a commercially available trace metals mixture (Trace) as well as a previously reported minimal trace metals solution (T2) (**Fig. 2D, Fig. S2A**).^36^ While we found that T2 decreased landomycins production, Trace increased it by ∼50 % when compared to the Co30 base medium.

To determine the ions responsible for inducing landomycin production in Trace, we evaluated the impact of six of the dominant ions in Trace (Zn^2+^, Co^2+^, Fe^3+^, Mn^2+^, Ni^2+^, and Cu^2+^) on landomycin titers (**Fig. 2D**). Addition of any one of the metals at their respective concentration in the trace mix was an improvement over the base medium, suggesting all six cations can activate the landomycin biosynthetic pathway, albeit to different extents. We found that Zn^2+^ was the major inducer of landomycin production in the trace mixture, achieving equivalent landomycin production at 10 µM as Co^2+^ did at 30 µM. Other individual ions yielded more modest landomycin titers. We then explored the additive effect of the metal ions by successively adding metal ions to the medium in an order based upon their individual effect. Addition of (i) Co^2+^, (ii) Co^2+^ and Fe^3+^, or (iii) Co^2+^, Fe^3+^, and Mn^2+^ to Zn^2+^ had no additional benefit relative to Zn^2+^ alone, while addition of Co^2+^, Fe^3+^, Mn^2+^, and Ni^2+^ yielded landomycins equivalent to the trace mixture. These observations suggest that a minimal solution of Zn^2+^, Co^2+^, Fe^3+^, Mn^2+^ and Ni^2+^ is sufficient to achieve landomycin yields equivalent to that of the complete Trace mix. Growth in 30 µM Zn^2+^, Mn^2+^, Ni^2+^, or Cu^2+^, did not improve yield further, nor did the addition of 30 µM Co^2+^, 20 µM Zn^2+^ or 30 µM Co^2+^ and 20 µM Zn^2+^ in combination with Trace (**Fig. S2B–C**). To validate that our improved production conditions are relevant at scale, we characterized the relative product yields of 1 L production cultures of Co30 and Trace. Trace produced more than double the total landomycin products compared to Co30 (915 vs. 430 µM). However, Trace produced less total and relative lanA, the pathway end product, compared to Co30 (223 vs 293 μM; 24 vs. 68% total products), and far more of the intermediates or byproducts lan, lanB, and A-lan (427 vs. 68 μM, 127 vs 49 µM, and 123 vs 12 µM, respectively).

**Figure 3:**
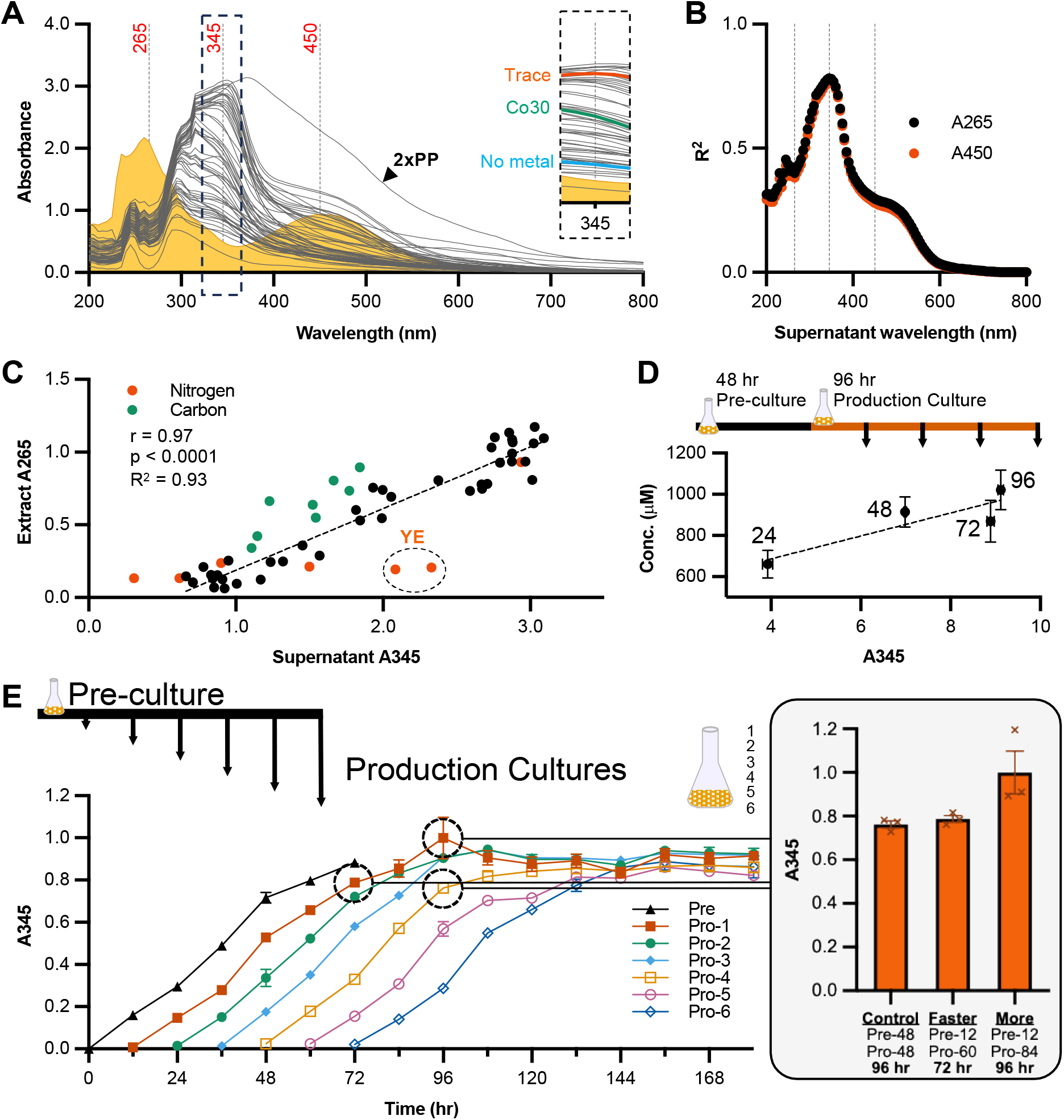
Detection of Relative Landomycin Production in Culture Supernatants. A) Average absorbance spectrum of the supernatant of 55 distinct production conditions tested for method optimization (grey lines). The absorbance spectrum of purified lanA (0.1 mg/mL) is shown in yellow. Vertical dashed lines indicate 265, 345, and 450 nm wavelengths. The spectrum of SG containing 2x phytone peptone (2xPP) is indicated by the label and black arrow. The inset depicts 345 nm with Trace, Co30 and No metal conditions labeled in orange, green, and blue, respectively. B) R^2^ values for linear regression analysis of supernatant absorbance at the given wavelength relative to the extract absorbance at 265 nm (black circles) or 450 nm (orange circles). Vertical dashed lines indicate 265 (R^2^ approx. 0.4), 345 (the maximum; R^2^ = 0.78), and 450 nm (R^2^ approx. 0.3) wavelengths. C) Plot of extract absorbance at 265 nm versus supernatant absorbance at 345 nm. Samples from media containing different nitrogen and carbon sources are orange and green, respectively. Yeast extract samples are distinguished by encircling and labeled YE. Nitrogen and carbon samples were removed from correlational and linear regression analysis due to apparent non-correlational clustering thought to result from the different base media absorbance. Pearson correlation coeff. and R^2^ of final 42 conditions are 0.965 (p<0.0001) and 0.93, respectively. D) Linear regression analysis of total landomycin concentration determined by LC-MS relative to A345 of culture time-course with samples taken every 24 hr from production cultures (R^2^ = 0.78). Arrows from production culture indicate sampling time points. E) Time-course analysis of pre-culture and production culture landomycin production determined by A345. Production cultures were set every 12 hr from the pre-culture and A345 was measured every 12 hrs (n=3). The inset depicts the magnitude of two different improved production methods (Faster and More), compared to the starting condition (Control), which decreased production time by 24 hr or increased yield by 32%.

### A rapid spectrophotometric approach to quantifying landomycins helps shorten production runs and improve yield

Although we were successful using A265 of culture extracts to improve landomycin production, the requirement of an extraction step limits sample throughput. Accordingly, we sought to develop a simplified method that directly used culture supernatant absorbance to determine relative landomycin titers. We assumed from the outset that the optimal wavelengths identified for sensitive detection of the purified compounds might be poor predictors in culture supernatants, due in part to the numerous contaminating compounds and altered chemical environment. To this end, we measured absorbance spectrum (200–800 nm) of culture supernatants for all 55 media compositions for which we had previously evaluated the absorbance of culture extracts while optimizing landomycin production. We plotted the average absorbance of the supernatants of each condition and found that the peaks identified in the purified landomycins or extracts (265 and 450 nm) were masked in culture supernatants (**Fig. 3A**). Accordingly, we ran linear regression analysis to generate trendlines for the average A265 of the 55 media conditions tested plotted versus the absorbance of the supernatant of the same 55 conditions taken at 5 nm intervals from 200–800 nm. Indeed, A265 and A450 of culture supernatants proved very poor predictors of landomycin titer (R^2^ = 0.4 & 0.3, respectively). To identify the most predictive wavelength, we plotted the R^2^ value for all trendlines calculated by correlating every supernatant absorbance wavelength with extract A265 (**Fig. 3B, 3A inset**). The maximum R^2^ was at 345 nm, with a value of 0.78, suggesting it is a good predictor of the extract A265, and thus total landomycin titer. When we plotted A265 of culture extracts versus A345 of culture supernatants, we were able to identify specific outlier sets within our data that likely resulted from different background media absorbances, including those autoclaved without glucose (Carbon) in the start media and those with different peptone sources (Nitrogen) (**Fig. 3C**). When we excluded these data from our analyses, the remaining 42 conditions retained a Pearson correlation coefficient of 0.97 (p<0.0001) and the trendline had a R^2^ of 0.93, both indicating A345 is an effective measure of total landomycin. We further validated the effectiveness of A345 of culture supernatants as a measure of landomycin production by directly measuring total landomycin production in production cultures by LC-MS at different timepoints during a production run (**Fig. 3D**). Finally, we the increased sample throughput permitted using supernatant A345 to optimize the landomycin production method by determining the optimal pre-culture subculturing timepoint and production culture endpoint (**Fig. 3E**). Starting at the pre-culture step, we subcultured pre-cultures into new production cultures every 12 hours, while monitoring landomycin production. We found that using a protocol that we named Faster, we could reduce the pre-culture from 48 hr to 12 hr, while extending the production culture from 48 hr to 60 hr, thereby reducing total production by 24 hr, without any loss in landomycin production. We also developed a protocol that we named More, where the production culture from a 12 hr pre-culture was extended to 84 hr, resulting in a further 32% increase in landomycin production compared to the starting condition (Control; 48 hr pre-culture, 48 hr production culture).

**Figure 4:**
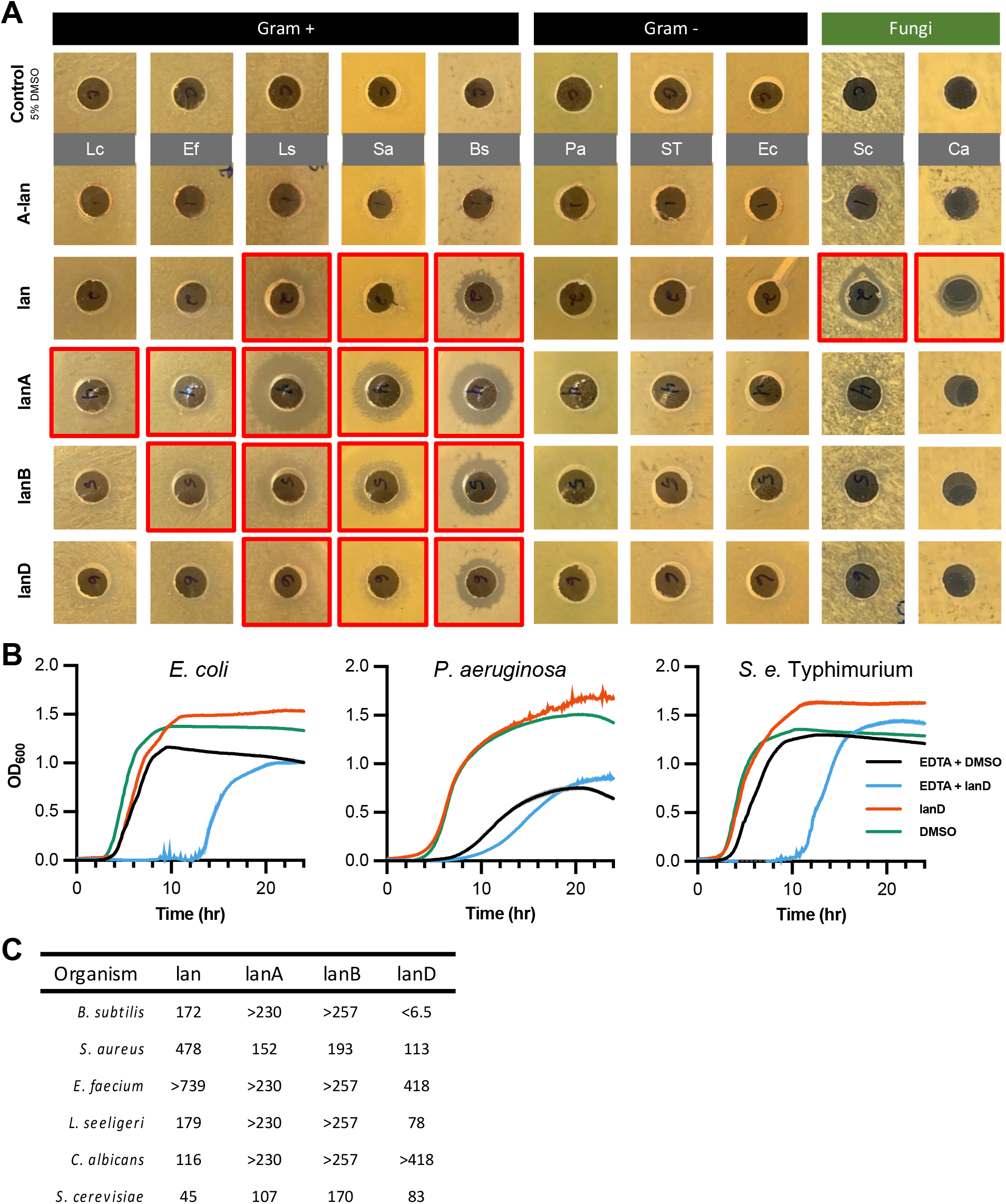
Antimicrobial spectrum of landomycin products of *S. cyanogenus* 136 pOOB92a. A) Inhibitory spectrum of anhydrolandomycinone (A-lan), landomycinone (lan), landomycin A (lanA), landomycin B (lanB), and landomycin D (lanD), against: *L. cremoris* (Lc), *E. faecium* (Ef), *L. seeligeri* (Ls), *S. aureus* (Sa), *B. subtilis* (Bs), *P. aeruginosa* (Pa), *S. enterica* ser. Typhimurium (ST), *E. coli* (Ec), *S. cerevisiae* (Sc), *and C. albicans* (Ca) as determined by agar well diffusion assay. 100 μL of 250 μg/mL of each compound was added to an agar well. Control contained the maximum DMSO concentration added to any well (5% v/v). Wells where inhibition of growth was visible are outlined with a bold red border. **B)** Growth curves of *E. coli*, *P. aeruginosa*, and *S.* Typhimurium in the presence of 250 ug/mL lanD with and without the addition of 1 mM EDTA. DMSO and EDTA+DMSO are provided as controls; growth inhibition by lanD required EDTA in Gram-negative organisms. **C)** Minimum Inhibitory concentration values (μM) calculated from growth inhibition curves (Fig. S5) for each active compound against organisms sensitive to at least one landomycin.

### Landomycins exhibit antibacterial and antifungal activities

Alhough landomycins are commonly reported to exert antimicrobial activity, the knowledge of their activity spectra are unknown.^38–40^ Additionally, the different landomycins, complex and impure mixtures, and/or qualitative metrics used in these studies limits comparison between them. We screened the five major landomycins found in production cultures for their antimicrobial activity against a diverse array of microbes (five Gram-positive, three Gram-negative, and two fungi) using an agar well diffusion assay. We found that lanA was active against all Gram-positive bacteria, while lan, lanB, and lanD inhibited at least three of the five Gram-positive strains tested (**Fig. 4A**). Lan also inhibited the growth of the fungi *S. cerevisiae* and *C. albicans*. A-lan was not active against any of the organisms tested, although this may be a result of the compound precipitating out when added to the medium. None of the landomycins inhibited the growth of Gram-negative bacteria. Given the broad antimicrobial activity we found landomycins had against Gram-positive bacteria and fungi, as well as the well-characterized anti-cancer activity many landomycins are known to exhibit, we postulated that the complete absence of growth inhibition seen with any Gram-negative bacterium may be due to the impermeability of the outer membrane. To investigate this hypothesis, we assessed whether permeabilization of the outer membrane using EDTA could sensitize Gram-negative bacteria to lanD.^41^ We initially determined non-inhibitory concentrations of EDTA for each Gram-negative strain (**Fig. S6**). The addition of 1 mM EDTA to growth medium was not inhibitory for *S.* Typhimurium or *E. coli*, but it was partially inhibitory to *P. aeruginosa*. As hypothesized, the addition of 1 mM EDTA sensitized both *S.* Typhimurium and *E. coli* to lanD, as evidenced by the delay in initiation of detectable growth from 4 h to 12 h (**Fig. 4B**). The growth delay for *P. aeruginosa* was less drastic due to the partial toxicity of EDTA. To better validate the bioactivity of the active landomycins, we also ran MIC assays on the landomycin-sensitive organisms *E. faecium*, *L. seeligeri*, *S. aureus*, *B. subtilis*, *S. cerevisiae*, and *C. albicans* (**Fig. S4**). Although we found these compounds frequently delayed growth, similar to our findings for lanD activity in the presence of EDTA for Gram-negative organisms, most compounds eventually permitted some microbial growth at the 24 h time point used to calculate the MIC (**Fig. S5 &** **Fig. 4C**). Per the MIC calculation, only lanD and lan completely inhibit the growth of *B. subtilis* and *S. cerevisiae*, respectively, at sub-50 µM concentrations. Overall, landomycins appear to have limited antimicrobial applications in their current form but show promise as starting scaffolds to synthesize a next generation of improved landomycins.

### Landomycins exhibit anticancer activity

We tested the efficacy of each of the landomycin products identified against human non-small cell lung cancer (NSCLC) cell line (A549-CCL-185; A549), using doxorubicin hydrochloride (DOX) as a positive control. After 24 hours of incubation with each compound of interest, we analyzed the A549 cells using flow cytometry. To determine the efficacy of each compound against A549 cells, we normalized to the negative control (0 µM), fit curves to each data set (**Fig. 5A**) and determined LC_50_ values for each drug of interest (**Fig. 5B**). These results indicate that while lanA is not as potent as DOX, it was the most potent member of the products of interest. For landomycins containing deoxyoligosaccharides, lanA was the most active while lanD was the least active. Additionally, we found that lan was one of the least active landomycins in this assay while A-lan was inactive up to 100 µM.

**Figure 5:**
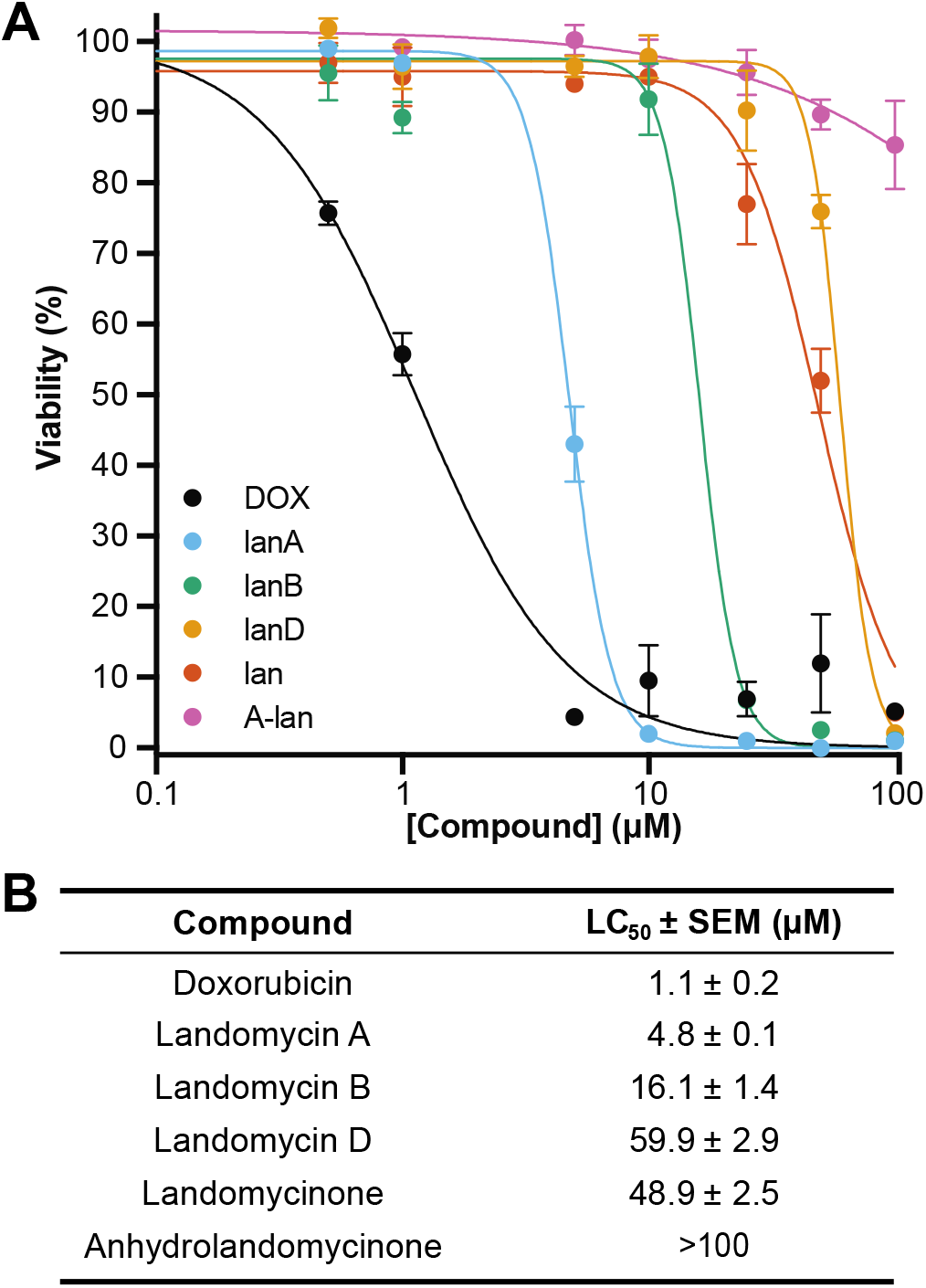
Viability of non-small cell lung carcinoma (A549) cells treated with landomycins. A) Dose response curves for landomycin products. We treated A549 cells with 0–100 µM DOX, lanA, lanB, lanD, lan, and A-lan for 24 hours prior to analysis via flow cytometry. We report the average percent viability (n=3), normalized to the 0 µM sample. Error bars reflecting the SEM. We fit each data point to a non-linear [inhibitor] versus response curve. B) tabulated LC50 values from dose response curves.

## Discussion

Unspecified production/induction conditions and low yield are major obstacles to the translation of natural products into viable therapeutics and chemical goods. Similarly, complicated separations add considerable costs and require the use of large amounts of energy and/or harmful solvents. Landomycins are not unique in this regard, and the lack of studies performed on this promising scaffold can be attributed, in part, to the difficulty in obtaining the molecule(s) of interest. To this end, we developed a spectrophotometric method using culture extracts to screen media compositions that reached lanA concentrations as high as 390 µM and a total polyketide concentration of 915 µM. By simplifying this method through the substitution of culture extracts with culture supernatants, we rapidly increased sample throughput, allowing us to monitor landomycin titers during production and across multiple cultures. Importantly, the underlying methodology that we developed, namely the identification of a single wavelength absorbed by all products of interest in their pure form and the correlation of this absorbance to an equally representative culture supernatant absorbance, can be easily extended to other natural products and their pathway intermediates that absorb light in the UV-vis range. Furthermore, by developing a differential centrifugation method to isolate landomycin-rich culture solids, we reduced the typically grueling purification method for lanA to a single normal phase silica column and precipitation step to yield >98% pure lanA.^10,34^ This application should also serve to simplify the isolation of other natural products expressed in culture solids of *Streptomyces*.

Of the landomycins investigated in this work (lanA, lanB, lanD, and lan), all compounds generally inhibited the growth of Gram-positive bacteria, lan inhibited the growth of fungi, and none of the compounds were active against Gram-negative bacteria without the addition of EDTA. LanA and lanB had the broadest spectrum of activity against Gram-positive bacteria in the agar well diffusion assay, but they generally had high MICs. Only lanD and lan had <50 μM MIC when treating *B. subtilis* and *S. cerevisiae,* respectively. Interestingly, lanA, lanB, and lanD only differ structurally by the length of the glycan chain, yet these subtle differences resulted in vastly different antimicrobial activity. In another study, landomycin R (lanR) – an anhydro-landomycin with a disaccharide glycan – was more inhibitory than landomycin Q, which only differs from lanR by an extra rhodinose on the glycan chain.^38^ Similarly, lanR only differs from lanD by the loss of an alcohol group and subsequent formation of a carbon-carbon double bond in the polyketide core, yet we found lanD to be far less active against *S. aureus* than was previously found for lanR (7 & 14 μM vs. 104 μM, respectively). Together, these findings better elucidate some of the activity-defining moieties of landomycins. In particular, the length and composition of the glycan chain are key to the bioactivity of landomycins and provide targets for future structure-function studies and synthetic modifications intended to improve potency and/or spectrum.

A common feature of many landomycins is their color (**Fig. 1A**), and during the course of the MIC assay, we noticed that the orange (lanA, lanB, lanD) or dark blue (lan) tinted media changed color to a darker, brown-black (**Fig. S7**). Even the aseptic media underwent this transformation. Given the underlying physical properties determinant of color, we hypothesize that a chemical reaction that alters the bonding in landomycin structures to be the likely cause. This suggests that the landomycins in this study are not particularly stable under the conditions tested, which could account for the transient growth inhibition found for many of the MIC assay samples. In fact, previous mechanistic investigations into the bioactivity of lanE noted both the rapid generation and subsequent decline of ROS in cancer cells, as well as the rapid generation and subsequent quenching of a fluorescent Michael adduct with biothiols, predicted to be mediated by the oxygen-dependent removal of the glycan.^11,13^ Taken with our findings, this suggests stabilization of the landomycin may be an effective strategy to generally improve landomycin bioactivity. Identification of the probable degradation products could further direct chemical modifications that improve landomycin stability and should be considered an important step in engineering a next generation of semi-synthetic landomycin therapeutics.

LanA and lanE are the most studied landomycins for their chemotherapeutic potential, which are reported to be mediated by inhibition of DNA synthesis, and/or oxidative stress to mitochondria via the rapid generation of reactive oxygen species (ROS).^11–13,42,43^ We found the LC_50_ of lanA to be 4.8 µM, which is within the range previously reported by the NCI for A549 cells (LC_50_ = 0.8 µM and ≥10 µM), and near the reported IC_50_.^44–47^ Concerning the broader activity trend of the glycated landomycins tested, we found an inverse relationship between oligosaccharide chain length and LD_50_ (lanA<lanB<lanD). This trend has been identified in other cell lines, which suggests that longer glycan chains generally enhance landomycin anticancer activity, and glycan composition can heavily impact activity.^32,34^ Though the aglycone lan was previously reported to have an IC_50_ below lanA for this cell line, we found lan to have reduced efficacy, and similar activity to lanD.^47^ Interestingly, by substituting the more common, but less informative, binary viability methods with a multiplexed, fluorescent dye assay, we identified that landomycin treated A549 cells undergo late-stage apoptosis, compared to DOX-treated cells that exhibit cellular necrosis (**Table S4**). Indeed, investigations into the mechanism of landomycin E bioactivity on Jurkat T-cell leukemia cells have noted that while both landomycins and DOX result in ROS generation, they may travel different paths to their common fate.^13,42,48^ Taken together, and given our improved production of the polyketide core, landomycins are promising chemotherapeutic candidates. Our results provide further guidance towards the designing and testing of new, semi-synthetic landomycins that better elucidate the relevant structural features by which landomycins mediate their cytotoxic effects, thereby improving candidate efficacy.

## Abbreviations

BGC: biosynthetic gene cluster
lanA: landomycin A
lanB: landomycin B
lanD: landomycin D
lan: landomycinone
A-lan: anhydrolandomycinone
Sc136: *S. cyanogenus* 136
Sc92a: *S. cyanogenus* 136 +pOOB92a
SG: soytone glucose medium
Co30: soytone glucose medium + 30 µM CoCl2
Trace: soytone glucose medium + 1× Trace metals solution

## Acknowledgments

We thank Prof. Bohdan Ostash (Ivan Franko Lviv National University, Lviv, Ukraine) for kindly providing us with *S. cyanogenus* 136 +pOOB92a. This work was supported by grants 1R21HD105934 (NIH) and CBET-2208390 (NSF) to N.U.N., grant CHE-2246963 (NSF) to C.S.B., and a generous gift from James Kanagy to C.R.M.

## References

1. Lee, N. et al. Mini review: Genome mining approaches for the identification of secondary metabolite biosynthetic gene clusters in Streptomyces. Comput. Struct. Biotechnol. J. 18, 1548–1556 (2020).

2. Liu, R., Yu, D., Deng, Z. & Liu, T. Harnessing in vitro platforms for natural product research: in vitro driven rational engineering and mining (iDREAM). Curr. Opin. Biotechnol. 69, 1–9 (2021).

3. Pham, J. V. et al. A review of the microbial production of bioactive natural products and biologics. Front. Microbiol. 10, 1–27 (2019).

4. Patridge, E., Gareiss, P., Kinch, M. S. & Hoyer, D. An analysis of FDA-approved drugs: Natural products and their derivatives. Drug Discov. Today 21, 204–207 (2016).

5. Harvey, A. L., Edrada-Ebel, R. & Quinn, R. J. The re-emergence of natural products for drug discovery in the genomics era. Nat. Rev. Drug Discov. 14, 111–129 (2015).

6. Li, C. J. & Trost, B. M. Green chemistry for chemical synthesis. Proc. Natl. Acad. Sci. U. S. A. 105, 13197–13202 (2008).

7. Chung, Y.-H. et al. Comparative Genomics Reveals a Remarkable Biosynthetic Potential of the Streptomyces Phylogenetic Lineage Associated with Rugose-Ornamented Spores . mSystems 6, 1–15 (2021).

8. Belknap, K. C., Park, C. J., Barth, B. M. & Andam, C. P. Genome mining of biosynthetic and chemotherapeutic gene clusters in Streptomyces bacteria. Sci. Rep. 10, 1–9 (2020).

9. Ward, A. C. & Allenby, N. E. Genome mining for the search and discovery of bioactive compounds: The Streptomyces paradigm. FEMS Microbiol. Lett. 365, 1–20 (2018).

10. Henkel, T., Rohr, J., Beale, J. M. & Schwenen, L. Landomycins, new angucycline antibiotics from streptomyces sp. I. Structural studies on landomycins A∼D. J. Antibiot. (Tokyo). 43, 492–503 (1990).

11. Terenzi, A. et al. Landomycins as glutathione-depleting agents and natural fluorescent probes for cellular Michael adduct-dependent quinone metabolism. Commun. Chem. 4, 1– 13 (2021).

12. Crow, R. T. et al. Landomycin A inhibits DNA synthesis and G1/S cell cycle progression. *Bioorganic Med*. Chem. Lett. 9, 1663–1666 (1999).

13. Panchuk, R. R. et al. Rapid generation of hydrogen peroxide contributes to the complex cell death induction by the angucycline antibiotic landomycin E. Free Radic. Biol. Med. 106, 134–147 (2017).

14. Westrich, L. et al. Cloning and characterization of a gene cluster from Streptomyces cyanogenus S136 probably involved in landomycin biosynthesis. FEMS Microbiol. Lett. 170, 381–387 (1999).

15. Matselyukh, B. P., Polishchuk, L. V, Lukyanchuk, V. V, Golembiovska, S. L. & Lavrenchuk, V. Y. Sequences of Landomycin E and Carotenoid Biosynthetic Gene Clusters, and Molecular Structure of Transcriptional Regulator of Streptomyces globisporus 1912. Mikrobiol. Z. 78, 60–70 (2016).

16. Feng, Z., Kallifidas, D. & Brady, S. F. Functional analysis of environmental DNA-derived type II polyketide synthases reveals structurally diverse secondary metabolites. Proc. Natl. Acad. Sci. U. S. A. 108, 12629–12634 (2011).

17. Peng, A. et al. Angucycline glycosides from an intertidal sediments strain Streptomyces sp. and their cytotoxic activity against hepatoma carcinoma cells. Mar. Drugs 16, (2018).

18. Kharel, M. K. et al. Angucyclines: Biosynthesis, mode-of-action, new natural products, and synthesis. Nat. Prod. Rep. 29, 264–325 (2012).

19. Weber, S., Zolke, C., Rohr, J. & Beale, J. M. Investigations of the biosynthesis and structural revision of landomycin A. J. Org. Chem. 59, 4211–4214 (1994).

20. Hrab, P. et al. Complete genome sequence of Streptomyces cyanogenus S136, producer of anticancer angucycline landomycin A. 3 Biotech 11, 1–10 (2021).

21. Bugaut, X., Guinchard, X. & Roulland, E. Synthesis of the Landomycinone Skeleton. J. Org. Chem. 75, 8190–8198 (2010).

22. Roush, W. R. & Neitz, R. J. Studies on the Synthesis of Landomycin A. Synthesis of the Originally Assigned Structure of the Aglycone, Landomycinone, and Revision of Structure. J. Org. Chem. 69, 4906–4912 (2004).

23. Guo, Y. & Sulikowski, G. A. Synthesis of the Hexasaccharide Fragment of Landomycin A: Application of Glycosyl Tetrazoles and Phosphites in the Synthesis of a Deoxyoligosaccharide. J. Am. Chem. Soc. 120, 1392–1397 (1998).

24. Yang, X., Fu, B. & Yu, B. Total synthesis of landomycin A, a potent antitumor angucycline antibiotic. J. Am. Chem. Soc. 133, 12433–12435 (2011).

25. Yalamanchili, S., Lloyd, D. & Bennett, C. S. Synthesis of the Hexasaccharide Fragment of Landomycin A Using a Mild, Reagent-Controlled Approach. Org. Lett. 21, 3674–3677 (2019).

26. Roush, W. R. & Bennett, C. E. A Highly Stereoselective Synthesis of the Landomycin A Hexasaccharide Unit. J. Am. Chem. Soc. 122, 6124–6125 (2000).

27. Yu, B. & Wang, P. Efficient Synthesis of the Hexasaccharide Fragment of Landomycin A: Using Phenyl 2,3-O-Thionocarbonyl-1-thioglycosides as 2-Deoxy-β-glycoside Precursors. Org. Lett. 4, 1919–1922 (2002).

28. Yang, X. & Yu, B. Synthesis of Landomycin D: Studies on the Saccharide Assembly. Synthesis (Stuttg). 48, 1693–1699 (2016).

29. Tanaka, H. et al. Combinatorial Synthesis of Deoxyhexasaccharides Related to the Landomycin A Sugar Moiety, Based on an Orthogonal Deprotection Strategy. Chem. – An Asian J. 5, 1407–1424 (2010).

30. Lee, J., Kang, S., Kim, J., Moon, D. & Rhee, Y. H. A Convergent Synthetic Strategy towards Oligosaccharides containing 2,3,6-Trideoxypyranoglycosides. Angew. Chemie – Int. Ed. 58, 628–631 (2019).

31. Von Mulert, U., Luzhetskyy, A., Hofmann, C., Mayer, A. & Bechthold, A. Expression of the landomycin biosynthetic gene cluster in a PKS mutant of Streptomyces fradiae is dependent on the coexpression of a putative transcriptional activator gene. FEMS Microbiol. Lett. 230, 91–97 (2004).

32. Shaaban, K. A., Stamatkin, C., Damodaran, C. & Rohr, J. 11-Deoxylandomycinone and landomycins X-Z, new cytotoxic angucyclin(on)es from a streptomyces cyanogenus K62 mutant strain. J. Antibiot. (Tokyo). 64, 141–150 (2011).

33. Luzhetskyy, A. et al. Generation of Novel Landomycins M and O through Targeted Gene Disruption. ChemBioChem 6, 675–678 (2005).

34. Shaaban, K. A., Srinivasan, S., Kumar, R., Damodaran, C. & Rohr, J. Landomycins P-W, cytotoxic angucyclines from streptomyces cyanogenus S-136. J. Nat. Prod. 74, 2–11 (2011).

35. Myronovskyi, M. et al. Generation of new compounds through unbalanced transcription of landomycin A cluster. Appl. Microbiol. Biotechnol. 100, 9175–9186 (2016).

36. Yushchuk, O. et al. Heterologous AdpA transcription factors enhance landomycin production in Streptomyces cyanogenus S136 under a broad range of growth conditions. Appl. Microbiol. Biotechnol. 102, 8419–8428 (2018).

37. Gessner, A. et al. Changing Biosynthetic Profiles by Expressing bldA in Streptomyces Strains. Chembiochem 16, 2244–2252 (2015).

38. Lai, Y. H. et al. Total synthesis of landomycins Q and R and related core structures for exploration of the cytotoxicity and antibacterial properties. RSC Adv. 11, 9426–9432 (2021).

39. Matseliukh, B. P., Konovalova, T. A., Polishchuk, L. V & Bambura, O. I. The sensitivity to landomycins A and E of streptomycetes, producers of polyketide antibiotics. Mikrobiol. Z. 60, 31–36 (1998).

40. Yushchuk, O. et al. Eliciting the silent lucensomycin biosynthetic pathway in Streptomyces cyanogenus S136 via manipulation of the global regulatory gene adpA. Sci. Rep. 11, 1–14 (2021).

41. Brown, M. R. W. & Richards, R. M. E. Effect of ethylenediamine tetraacetate on the resistance of pseudomonas aeruginosa to antibacterial agents [17]. Nature 207, 1391–1393 (1965).

42. A, K., et al. Mechanisms underlying the anticancer activities of the angucycline landomycin E. Biochem. Pharmacol. 74, 1713–1726 (2007).

43. Lehka, L. V., Panchuk, R. R., Berger, W., Rohr, J. & Stoika, R. S. The role of reactive oxygen species in tumor cells apoptosis induced by landomycin A. Ukr. Biochem. J. 87, 72–82 (2015).

44. Shoemaker, R. H. The NCI60 human tumour cell line anticancer drug screen. Nat. Rev. Cancer 6, 813–823 (2006).

45. Shankavaram, U. T. et al. CellMiner: A relational database and query tool for the NCI-60 cancer cell lines. BMC Genomics 10, 1–10 (2009).

46. Reinhold, W. C. et al. CellMiner: A Web-Based Suite of Genomic and Pharmacologic Tools to Explore Transcript and Drug Patterns in the NCI-60 Cell Line Set. Cancer Res. 72, 3499–3511 (2012).

47. Zhang, Y. et al. Himalaquinones A–G, Angucyclinone-Derived Metabolites Produced by the Himalayan Isolate Streptomyces sp. PU-MM59. J. Nat. Prod. **84**, 1930–1940 (2021).

48. Panchuk, R. R. Signaling pathways involved in apoptosis induced by novel angucycline antibiotic landomycin E in Jurkat T-Leukemia cells. 27, 124–131 (2011).

