## Supplemental Information for "Rapid spectrophotometric detection for optimized production of landomycins and characterization of their therapeutic potential"

### Materials and Methods

#### Microbial MIC calculation

The MIC for each compound that exhibited complete inhibition was calculated by plotting the log of the concentration versus the $OD_{600}$ normalized to control condition, and then performing nonlinear regression (curve fit) in GraphPad Prism 9.5.0 using the equations[1]:

$$\text{Eq 1: } \mathrm{M} = \mathrm{logMIC} - \frac{1}{\mathrm{B}}$$

$$\text{Eq 2: } Y = \mathrm{A} + \mathrm{C} \times e^{\left(-e^{(\mathrm{B}(\mathrm{X}-\mathrm{M}))}\right)}$$

Where M is the log concentration of the inflexion point, logMIC is the log value of the MIC, B is the slope, A is the lower asymptote of y (approx. 0), and C is the distance between the lower and upper asymptote (approx. 1). The rules for the initial values for the variables were: logMIC was set to the value of X at YMID, B was selected as the initial value to be fit, A was set to the YMIN (ideally ~0), and C was set to YMAX-YMIN (the range). For negative OD values, OD were changed to 0. For compounds where complete growth inhibition was not reached, the MIC was determined to be greater than the maximum value tested. For compounds where no growth was visible even at the lowest concentrations, MIC was determined to be less than the minimum value tested.

## LC-MS

Culture extracts were analyzed for landomycins using targeted LC-MS experiments on a quadrupole-time of flight instrument (AB Sciex, TripleTOF 5600+). Chromatographic separation was performed on a C18 column (Synergi 4 µm Hydro-RP 80 Å, 250 x 2 mm, Phenomenex) as gradient indicated at Table S5. Solvent A was 0.1% formic acid in water (v/v). Solvent B was 0.1% formic acid in methanol (v/v). Injection volume was 10 µL, and the oven temperature was set to 15°C. The flow rate was kept constant at 0.2ml/min. Product Ion scan experiments were set for each compound of interest as Table S6. Areas under curve (AUC) were interpolated using Skyline (22.2.0.255) and normalized with total ion currents (TIC). Each compound's corresponding product ions used for quantification and retention times were also included in Table S6.

**Figures**

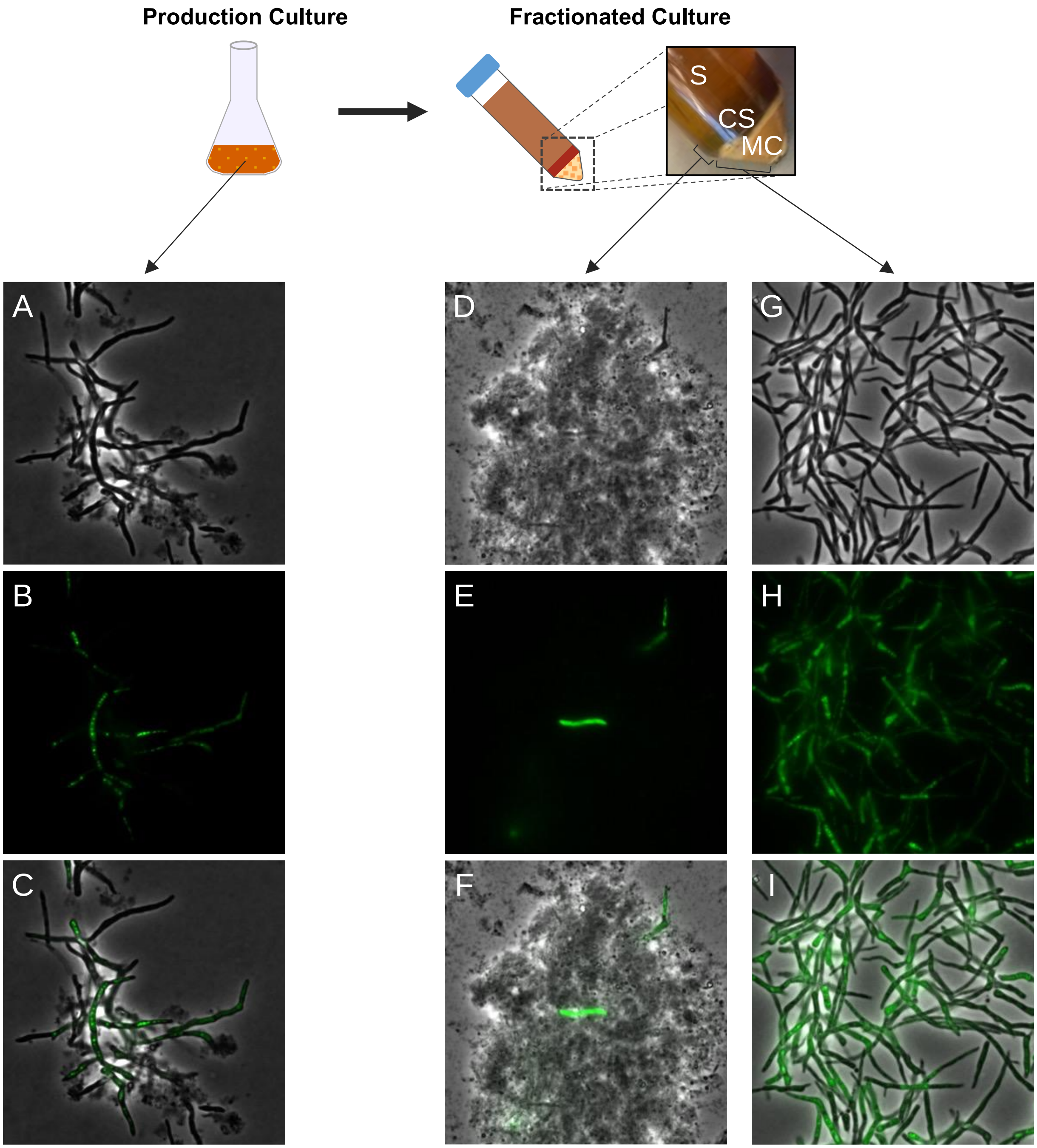


**Figure S1: Morphology of production cultures, culture solids, and mycelia/cell fractions.** Phase contrast (A,D,G) fluorescence (B,E,H), and overlay (C, F, I) microscopy images of production cultures (A-C) and fractionated Culture solids (CS; D-F) and mycelia/cells (MC; G-I). Samples were stained with SYBRSafe DNA stain prior to imaging to facilitate identification of intact cells and were imaged at 100x magnification.

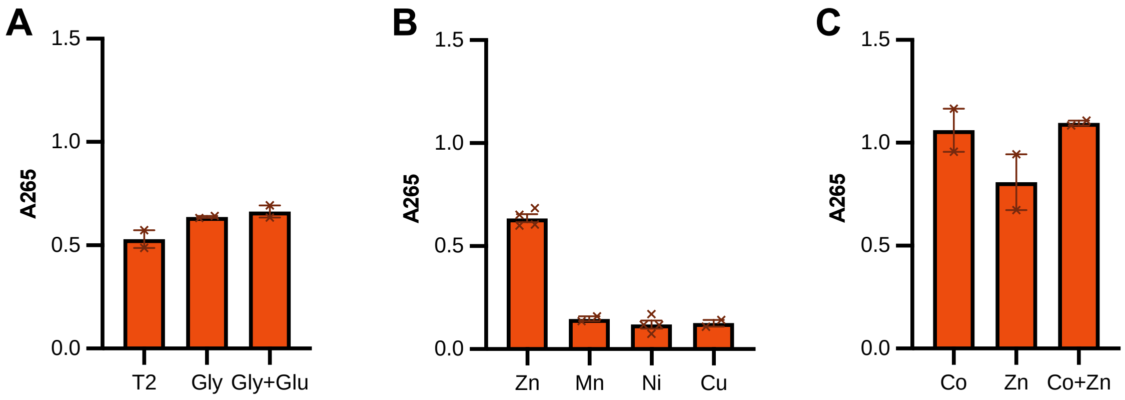


**Figure S2: Relative Landomycin Production in Additional Production Medium Compositions.** A) Landomycin production in SG with A) Trace 2 metals (T2), 2% glycerol (Gly) or 1% glycerol and 1% glucose (Gly+Glu); B) 30 µM Zn, Mn, Ni, or Cu; C) Trace with 30 µM Co, 20 Zn, or 30 µM Co + 20 µM Zn.

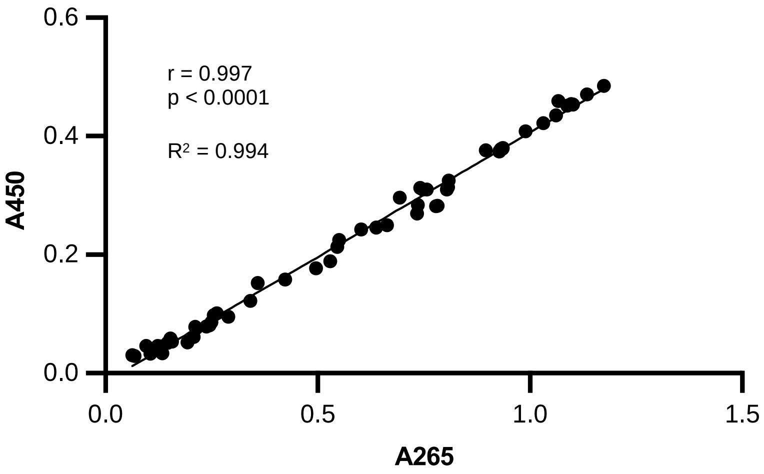


**Figure S3: Correlational Analysis of Extract A265 and A450.** Extract absorbance at 265 nm plotted vs. extract absorbance at 450 nm. Pearson correlation coefficient is 0.997 ($p<0.0001$), indicating a strong correlation between absorbance measurements at these wavelengths.

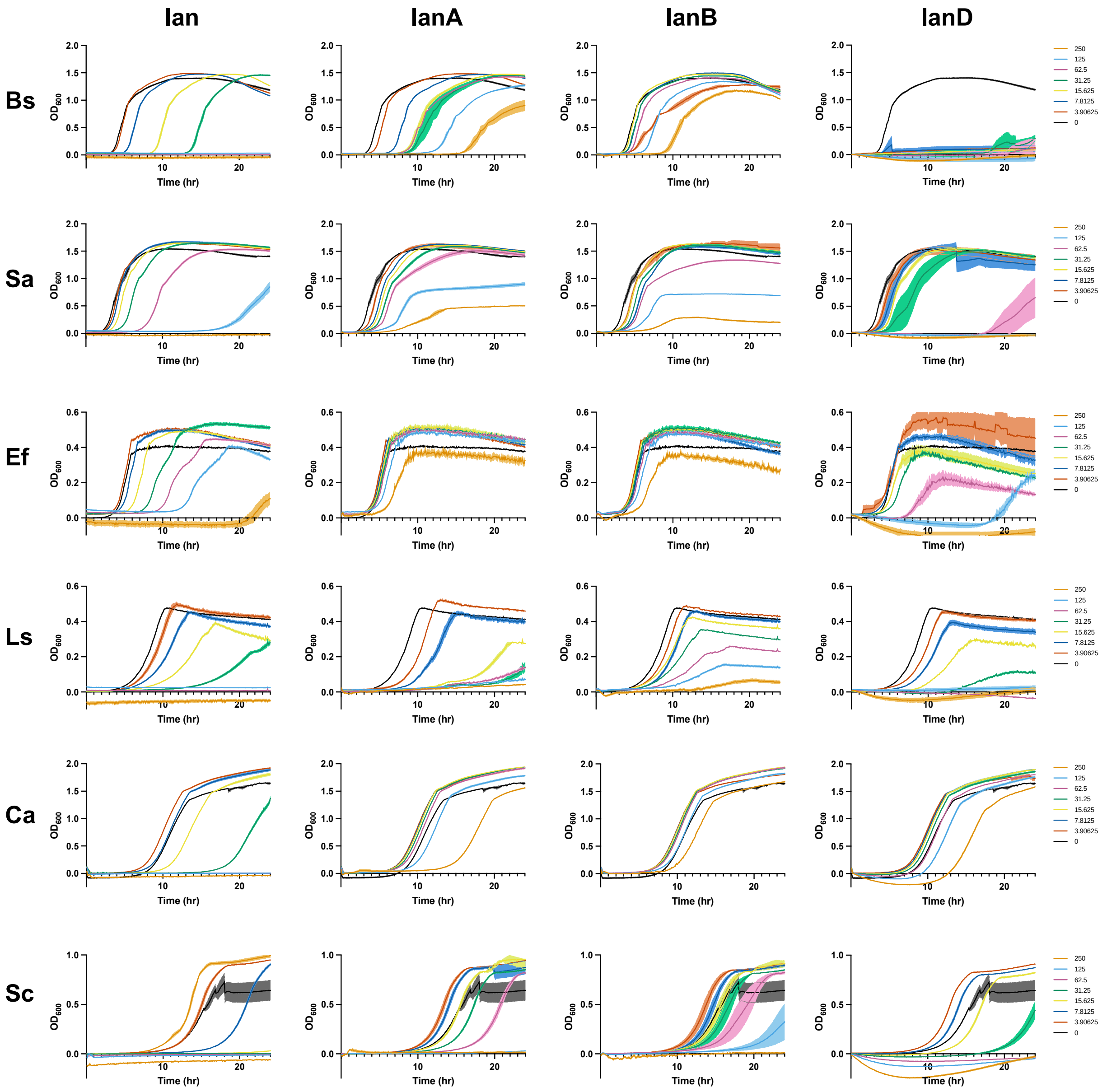


**Figure S4: Growth curves used for MIC determination.** Growth curves of *B. subtilis* (Bs), *S. aureus* (Sa), *E. faecium* (Ef), *L. seeligeri* (Ls), *C. albicans* (Ca), and *S. cerevisiae* (Sc) in the presence of different concentrations of landomycinone (lan), landomycin A (lanA), landomycin B (lanB), and landomycin D (lanD).

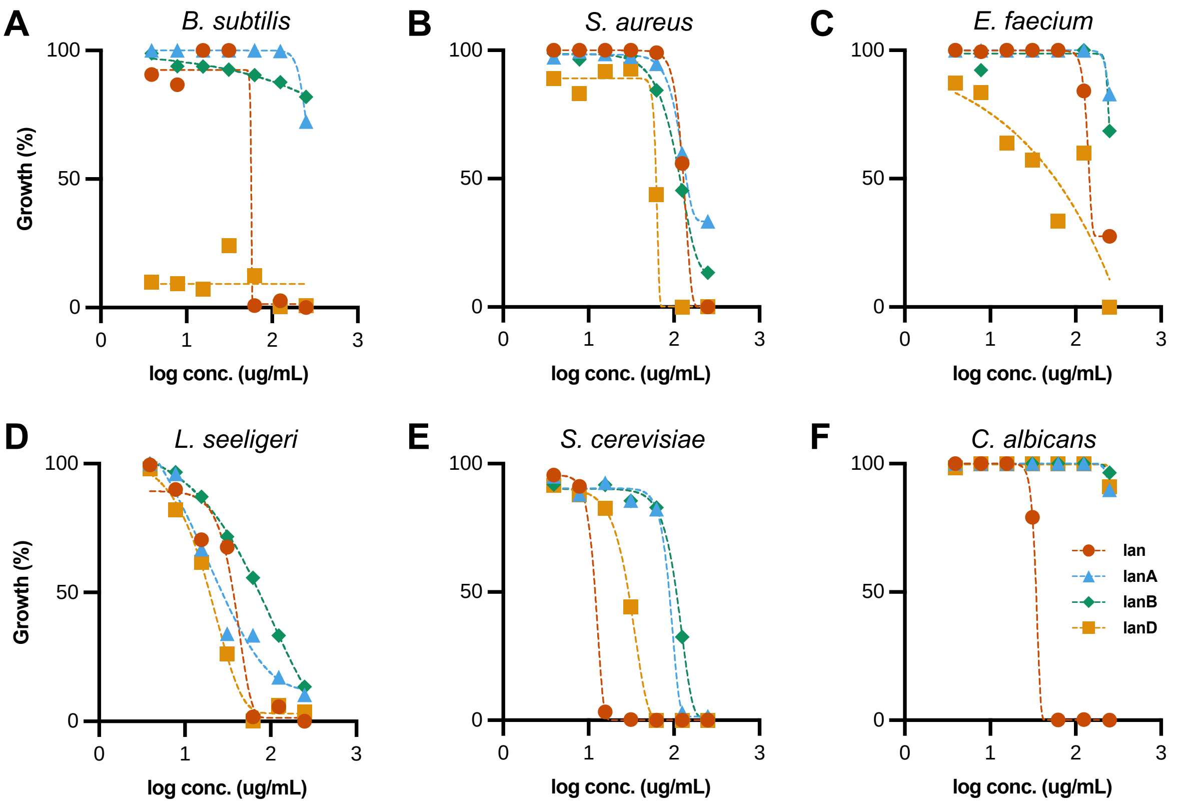


**Figure S5: Minimum Inhibitory Concentration of Landomycins.** Growth inhibition plots of landomyninone (lan; orange circles), landomycin A (lanA; blue triangles), landomycin B (lanB; green diamonds), and landomycin D (lanD; yellow squares) calculated at 24 hrs for A) *B. subtilis*, B) *S. aureus*, C) *E. faecium*, D) *L. seeligeri*, E) *S. cerevisiae*, and F) *C. albicans*. Non-linear regression was performed to fit curves to growth inhibition data (color matched to symbol color) and calculate MIC values (Fig. 4B).

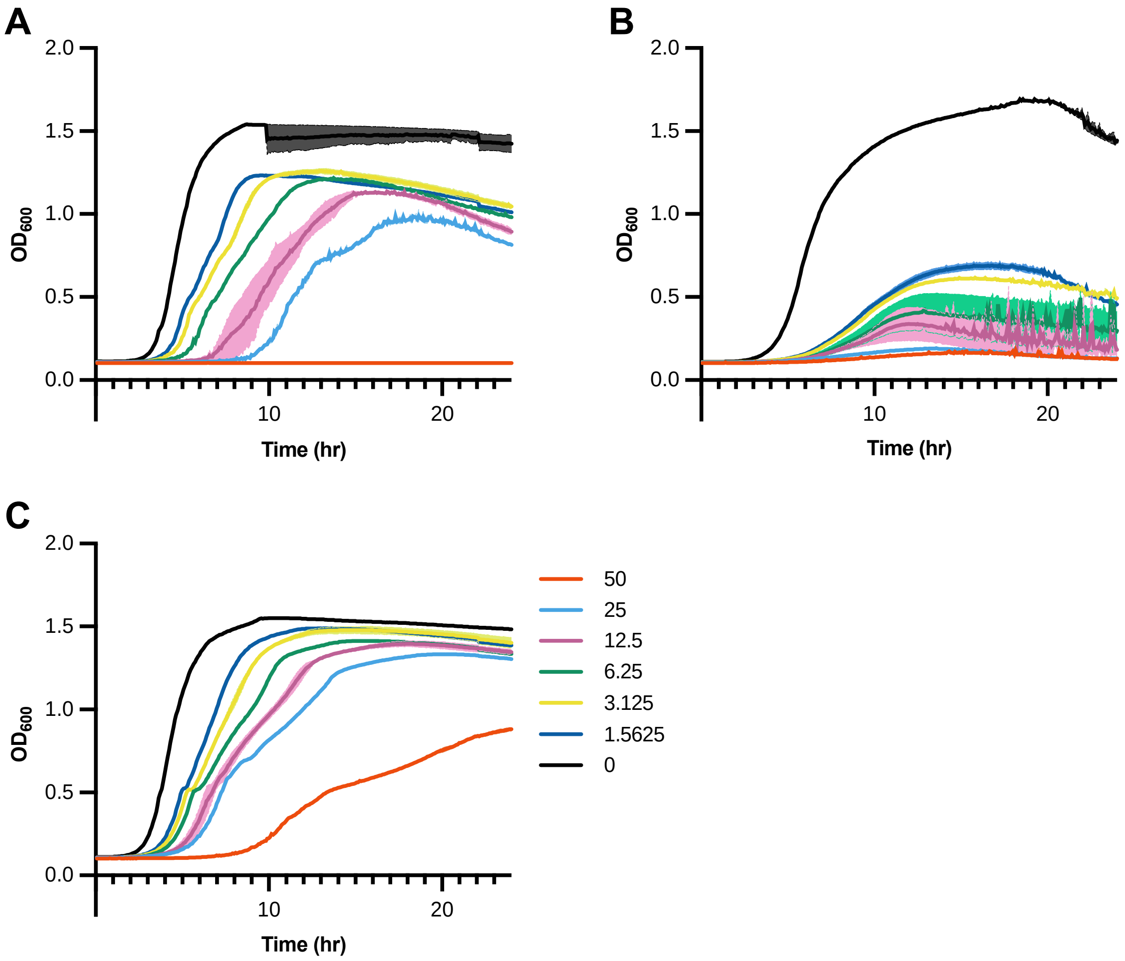


**Figure S6: Gram-negative EDTA growth inhibition.** Growth curves of A) *E. coli*, B) *P. aeruginosa*, and C) *S. typhimurium* grown in BHI containing different concentrations of EDTA (0-50 mM)**.**

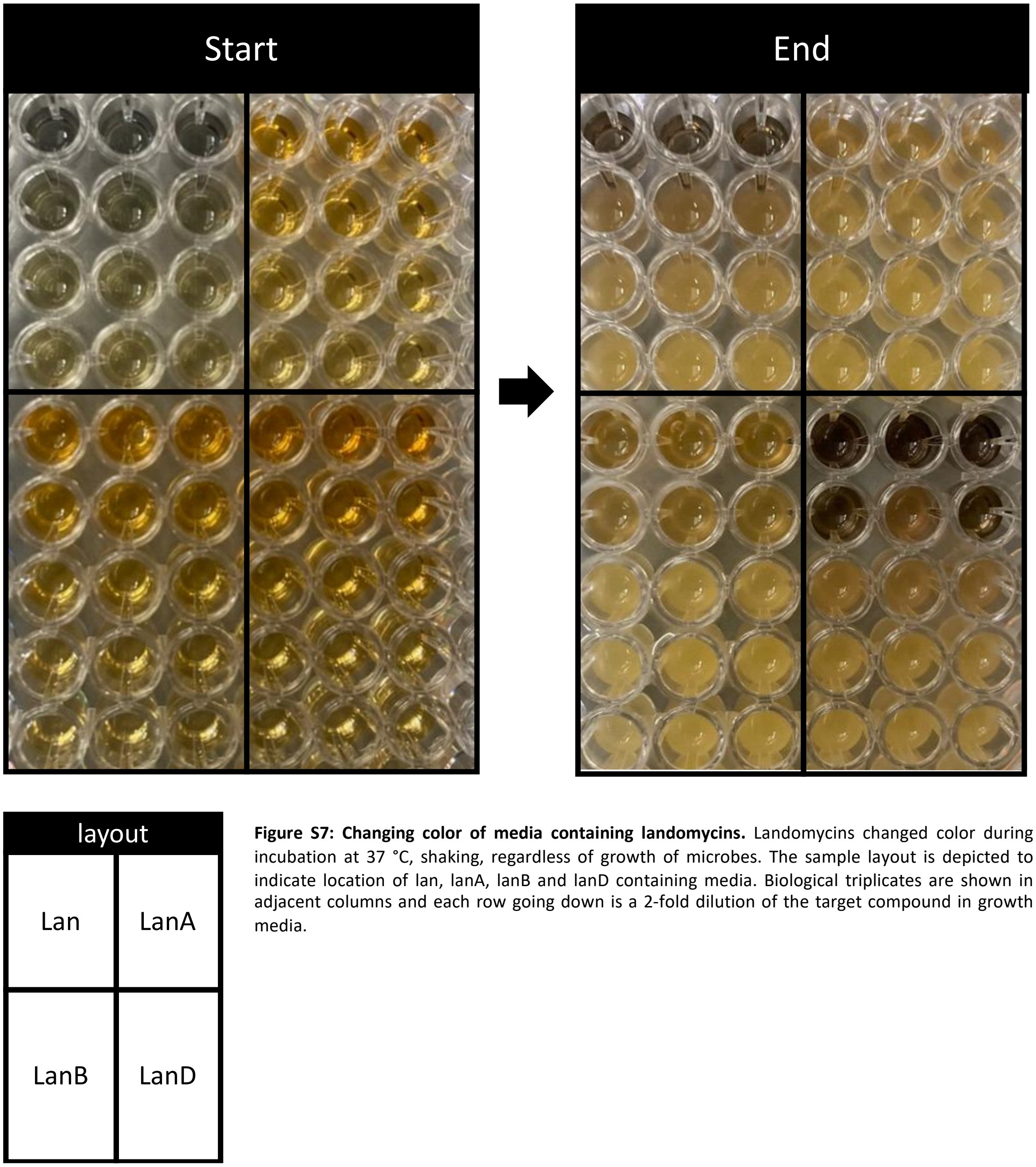


**Figure S7: Changing color of media containing landomycins.** Landomycins changed color during incubation at 37 °C, shaking, regardless of growth of microbes. The sample layout is depicted to indicate location of lan, lanA, lanB and lanD containing media. Biological triplicates are shown in adjacent columns and each row going down is a 2-fold dilution of the target compound in growth media.

### Tables

**Table S1**: Strains used in this study.

| **Strain** | **Genotype/Use/Origin** | **Source/Origin** |
|---|---|---|
| *Streptomyces cyanogenus* 136 +pOOB92a (Sc92a) | *Streptomyces cyanogenus* 136 with plasmid pOOB92a integrated into the genome. | Dr. Bohdan Ostash, Lviv University |
| *Pseudomonas aeruginosa* | Strain PA14. Burn wound isolate. | Dr. Roberto Kolter, HMS |
| *Staphylococcus aureus* | Strain UAMS-1. Osteomyelitis isolate. | Dr. Abraham L. Sonenshein, TUSM |
| *Escherichia coli* Nissle 1917 | Probiotic strain. Human fecal isolate. | iGEM team, Tufts University |
| *Salmonella enterica* subsp. *enterica* serovar Typhimurium | Strain SL1344. Auxotrophic mutant of S2337 (also known as 4/74 and S2337/65), an isolate from a calf with salmonellosis. | Dr. Dayong Wu, Tufts U. |
| *Lactococcus cremoris* | Subsp. c*remoris, s*train MG1363. | Dr. Jan Peter van Pijkeren, UW |
| *Enterococcus faecium* | Strain NRRL B-2354. | NRRL |
| *Listeria seeligiri* | Strain NRRL B-33019. | NRRL |
| *Bacillus subtilis* | Strain 168. | BGSC |
| *Saccharomyces cereviseae* | Strain BMA64-1A. | Euroscarf |
| *Candida albicans* | Strain SC5314. | Dr. Carol Kumamoto, TUSM |

**Table S2. Filter/Laser Combinations for Viability Dyes**

| Cell Stain | Annexin V-FITC | Ethidium homodimer I | Hoechst 33342 |
|---|---|---|---|
| Ex/Em | 485/535 | 528/617 | 350/461 |
| Filter - Laser | Green – B | Red – B | Blue – V |
| Emission Filter | 512/18 | 695/50 | 450/45 |

**Table S3. Compensation Control Values to Remove Spectral Overlap**

| | Compensation Value | | | |
|---|---|---|---|---|
| **Channel of Interest** | **GRN-B** | **RED-B** | **BLU-V** | **YEL-B** |
| GRN-B | -- | -- | 5.0 | -- |
| RED-B | 9.5 | -- | -- | 11.5 |
| BLU-V | -- | -- | -- | -- |
| YEL-B | -- | 45.7 | -- | -- |

*“--" denotes no compensation needed*

**Table S4.** Raw viable, apoptotic, and necrotic A549 cell populations under challenge conditions

| Compound | [Compound] (μM) | Average Viable (%) | Viable SEM | Average Early Apoptotic (%) | Early Apoptotic SEM | Average Late Apoptotic (%) | Late Apoptotic SEM | Average Necrotic (%) | Necrotic SEM |
|---|---|---|---|---|---|---|---|---|---|
| DOX | 0 | 89.9 | 0.8 | 2.2 | 0.2 | 4.6 | 0.6 | 3.3 | 0.2 |
| | 0.5 | 68.1 | 1.6 | 3.9 | 0.3 | 12.3 | 1.4 | 15.7 | 0.5 |
| | 1 | 50.1 | 3.0 | 5.0 | 0.8 | 16.1 | 1.6 | 28.7 | 1.1 |
| | 5 | 3.9 | 0.7 | 0.3 | 0.1 | 25.3 | 2.1 | 70.4 | 1.5 |
| | 10 | 8.5 | 5.1 | 0.5 | 0.1 | 26.9 | 4.5 | 64.1 | 9.4 |
| | 25 | 6.2 | 2.4 | 0.4 | 0.1 | 18.8 | 1.3 | 74.5 | 3.3 |
| | 50 | 10.8 | 7.0 | 0.6 | 0.4 | 17.4 | 5.6 | 71.2 | 12.9 |
| | 100 | 5.2 | 0.8 | 0.0 | 0.0 | 20.6 | 4.9 | 74.2 | 21.4 |
| lanA | 0 | 92.8 | 0.7 | 1.1 | 0.2 | 3.7 | 0.3 | 2.3 | 0.3 |
| | 0.5 | 91.0 | 0.3 | 1.4 | 0.2 | 4.7 | 0.3 | 2.9 | 0.1 |
| | 1 | 88.9 | 1.1 | 1.8 | 0.3 | 6.5 | 0.4 | 2.7 | 0.6 |
| | 5 | 39.3 | 5.3 | 2.8 | 0.5 | 41.5 | 3.5 | 16.5 | 1.8 |
| | 10 | 1.9 | 0.4 | 0.5 | 0.1 | 80.0 | 1.6 | 17.6 | 1.6 |
| | 25 | 1.4 | 0.8 | 1.0 | 0.5 | 95.9 | 0.8 | 1.7 | 0.6 |
| | 50 | 0.0 | 0.0 | 0.6 | 0.3 | 98.2 | 0.5 | 1.2 | 0.3 |
| | 100 | 0.5 | 0.2 | 1.2 | 0.2 | 97.0 | 0.8 | 1.4 | 0.6 |
| lanB | 0 | 82.3 | 4.7 | 13.7 | 3.5 | 3.0 | 0.9 | 1.0 | 0.4 |
| | 0.5 | 78.7 | 3.9 | 16.2 | 2.8 | 4.1 | 1.1 | 1.0 | 0.2 |
| | 1 | 73.4 | 2.2 | 20.9 | 1.5 | 4.3 | 1.0 | 1.3 | 0.3 |
| | 5 | 87.0 | 1.8 | 10.0 | 1.4 | 2.0 | 0.3 | 1.0 | 0.2 |
| | 10 | 75.6 | 5.1 | 14.1 | 3.4 | 8.9 | 1.4 | 1.4 | 0.3 |
| | 25 | 5.6 | 0.8 | 31.9 | 4.1 | 59.9 | 3.3 | 2.7 | 0.7 |
| | 50 | 2.1 | 0.5 | 90.1 | 2.4 | 5.5 | 0.9 | 2.3 | 0.9 |
| | 100 | 0.9 | 0.2 | 82.1 | 2.7 | 16.0 | 2.7 | 0.9 | 0.2 |
| lanD | 0 | 0.9 | 0.2 | 82.1 | 2.7 | 16.0 | 2.7 | 0.9 | 0.2 |
| | 0.5 | 88.0 | 1.4 | 6.2 | 0.3 | 4.6 | 0.9 | 1.2 | 0.2 |
| | 1 | 83.2 | 3.1 | 9.6 | 1.5 | 5.0 | 1.3 | 2.1 | 0.4 |
| | 5 | 83.3 | 1.5 | 10.0 | 0.9 | 4.9 | 0.6 | 1.8 | 0.2 |
| | 10 | 84.5 | 3.0 | 8.2 | 2.0 | 5.5 | 1.0 | 1.8 | 0.1 |
| | 25 | 77.9 | 5.7 | 12.7 | 2.8 | 7.6 | 2.6 | 1.7 | 0.4 |
| | 50 | 65.6 | 2.4 | 16.9 | 1.5 | 15.0 | 1.0 | 2.6 | 0.3 |
| | 100 | 1.8 | 0.3 | 46.5 | 1.9 | 43.9 | 2.7 | 7.8 | 1.3 |
| lan | 0 | 88.7 | 2.0 | 5.8 | 0.7 | 3.9 | 1.2 | 1.6 | 0.2 |
| | 0.5 | 86.0 | 2.8 | 6.6 | 0.8 | 5.6 | 1.8 | 1.7 | 0.2 |
| | 1 | 84.4 | 4.1 | 8.4 | 2.3 | 5.1 | 1.3 | 2.0 | 0.6 |
| | 5 | 83.2 | 0.5 | 7.9 | 0.4 | 6.9 | 0.6 | 2.0 | 0.1 |
| | 10 | 84.6 | 0.8 | 8.3 | 0.5 | 5.0 | 0.4 | 2.2 | 0.1 |
| | 25 | 68.7 | 5.7 | 13.7 | 1.9 | 15.1 | 3.7 | 2.6 | 0.3 |
| | 50 | 46.0 | 4.5 | 15.4 | 0.9 | 34.9 | 3.1 | 3.8 | 0.6 |
| | 100 | 4.3 | 1.1 | 35.2 | 1.2 | 54.1 | 1.2 | 6.4 | 1.2 |
| A-lan | 0 | 86.0 | 0.9 | 9.7 | 1.0 | 3.2 | 0.2 | 1.1 | 0.2 |
| | 0.5 | 89.7 | 1.1 | 6.2 | 1.1 | 3.0 | 0.5 | 1.1 | 0.2 |
| | 1 | 85.3 | 0.9 | 8.9 | 1.2 | 4.4 | 0.4 | 1.4 | 0.1 |
| | 5 | 86.2 | 2.1 | 6.8 | 0.9 | 5.0 | 1.4 | 1.9 | 0.2 |
| | 10 | 83.9 | 2.8 | 9.0 | 1.9 | 5.4 | 1.0 | 1.7 | 0.1 |
| | 25 | 82.3 | 3.2 | 10.6 | 1.9 | 5.4 | 1.5 | 1.7 | 0.3 |
| | 50 | 77.1 | 2.1 | 11.8 | 1.2 | 8.6 | 1.2 | 2.5 | 0.5 |
| | 100 | 73.4 | 6.3 | 11.7 | 1.4 | 12.3 | 4.8 | 2.5 | 0.3 |

**Table S5: LC-MS Gradient method for landomycins.**

| Time (min) | Flow Rate (mL/min) | % Solvent A | % Solvent B |
|---|---|---|---|
| 0 | 0.2 | 97 | 3 |
| 8 | 0.2 | 97 | 3 |
| 38 | 0.2 | 5 | 95 |
| 45 | 0.2 | 5 | 95 |
| 47 | 0.2 | 97 | 3 |
| 65 | 0.2 | 97 | 3 |

**Table S6: MS and Quantification parameters for landomycins.**

| Landomycins | Polarity | Declustering Potential | Collision Energy | Retention Time | Precursor(m/z) | Product (m/z) |
|---|---|---|---|---|---|---|
| Landomycinone | Negative | 65 | 35 | 42.54 | 337.07 | 318.05 |
| Landomycin A | Negative | 30 | 70 | 42.97 | 1085.46 | 925.41 |
| Landomycin B | Negative | 30 | 60 | 42.05 | 971.39 | 811.34 |
| Landomycin D | Negative | 30 | 40 | 40.9 | 597.20 | 437.15 |

#### Structure Characterization

Landomycin A

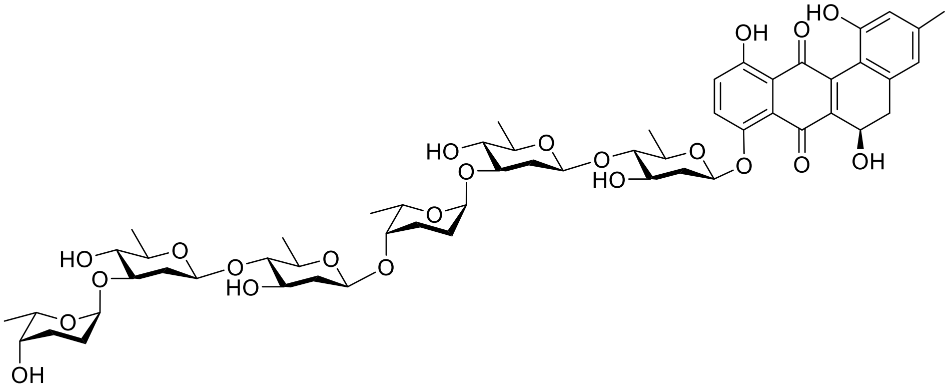

Chemical Formula: $C_{55}H_{74}O_{22}$

$^1$H NMR (500 MHz, $CDCl_3$) δ 12.35 (s, 1H), 8.87 (s, 1H), 7.55 (d, *J* = 9.4 Hz, 1H), 7.28 (s, 1H), 6.80 (s, 1H), 6.76 (s, 1H), 5.08 (dd, *J* = 8.9, 3.0 Hz, 2H), 4.96 (s, 2H), 4.70 (s, 1H), 4.62 (s, 1H), 4.56 – 4.43 (m, 4H), 4.34 (s, 1H), 4.18 – 4.03 (m, 2H), 3.72 (m, 1H), 3.65 (s, 1H), 3.60 (dq, *J* = 13.0, 6.2 Hz, 1H), 3.56 – 3.47 (m, 3H), 3.39 (td, *J* = 8.0, 5.3 Hz, 3H), 3.29 (dt, *J* = 12.4, 6.2 Hz, 1H), 3.15 – 3.02 (m, 4H), 2.98 (t, *J* = 8.8 Hz, 1H), 2.90 (dd, *J* = 15.8, 4.5 Hz, 1H), 2.80 (d, *J* = 4.4 Hz, 1H), 2.73 (m, 1H), 2.32 (m, 4H), 2.28 – 2.19 (m, 2H), 2.14 (m, 1H), 2.07 – 1.87 (m, 5H), 1.77 (d, *J* = 11.2 Hz, 1H), 1.73 – 1.61 (m, 4H), 1.39 (dd, *J* = 8.0, 6.1 Hz, 6H), 1.27 (dd, *J* = 16.5, 6.2 Hz, 6H), 1.22 (dd, *J* = 8.6, 6.5 Hz, 6H).

$^{13}$C NMR (126 MHz, $CDCl_3$) δ 192.74, 182.95, 159.75, 155.14, 150.71, 146.93, 143.71, 138.76, 136.69, 132.82, 126.75, 123.71, 120.14, 119.25, 114.89, 113.26, 101.48, 100.90, 100.86, 99.69, 97.79, 97.50, 88.52, 87.87, 80.56, 80.37, 75.77, 75.26, 75.23, 72.35, 72.26, 70.86, 70.34, 69.64, 69.24, 67.72, 67.09, 62.38, 38.27, 37.62, 37.13, 37.02, 36.29, 25.48, 25.07, 24.46, 24.09, 21.19, 17.84, 17.82, 17.80, 17.01, 16.94.

ESI-MS: [M+Na]: 1109.45

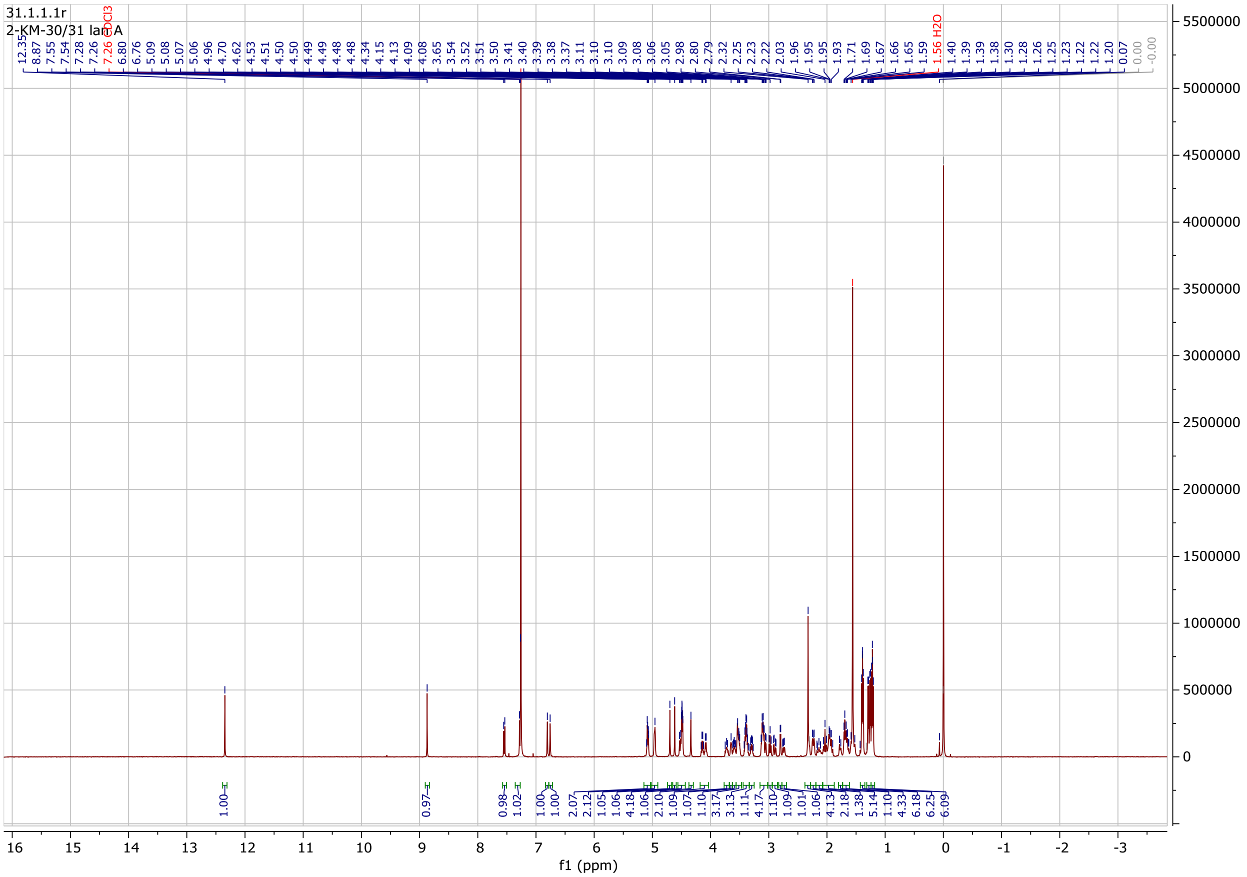

31.1.1.1r
2-KM-30/31 lam A
f1 (ppm)

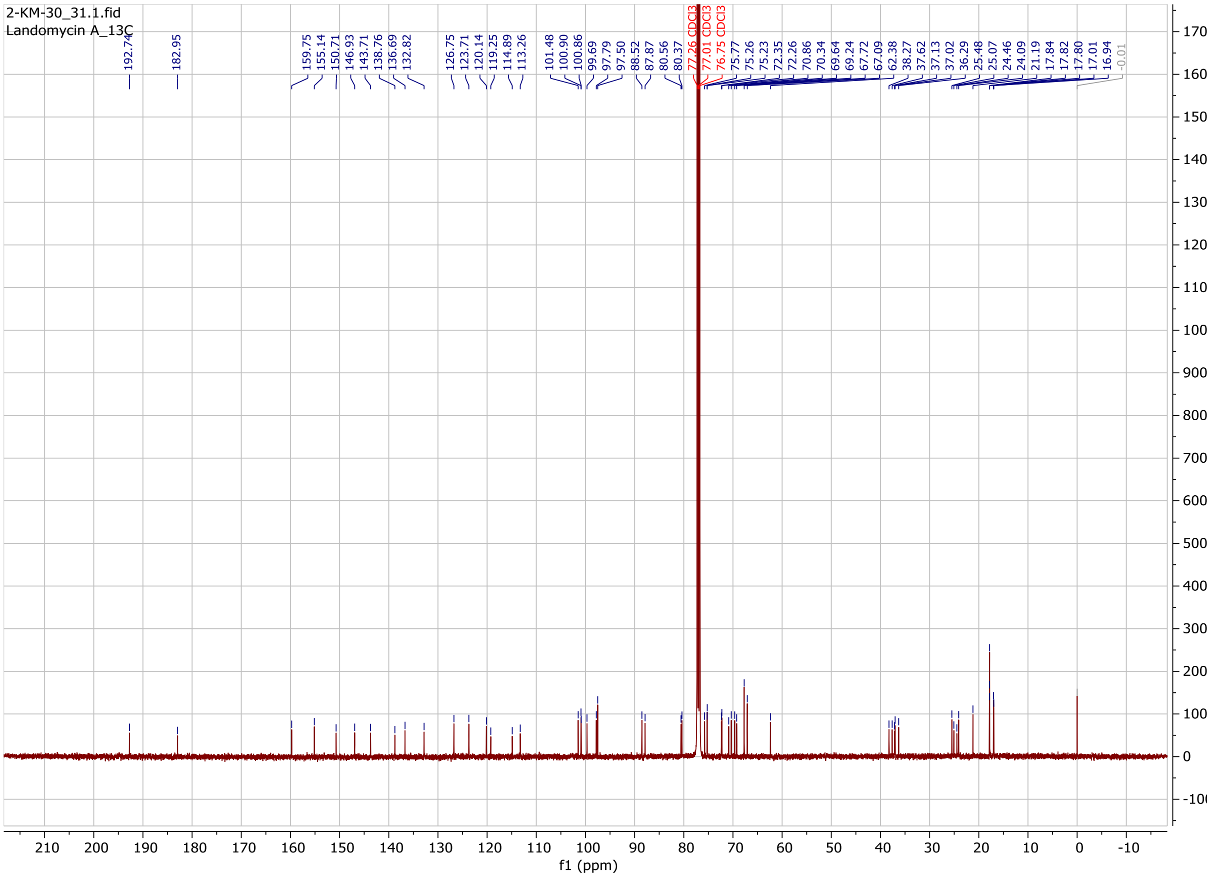
2-KM-30_31.1.fid
Landomycin A_13C
192.74
182.95
159.75
155.14
150.71
146.93
143.71
138.76
136.69
132.82
126.75
123.71
120.14
119.25
114.89
113.26
101.48
100.90
100.86
99.69
97.79
97.50
88.52
87.87
80.56
80.37
77.26 CDCl3
77.01 CDCl3
76.75 CDCl3
75.77
75.26
75.23
72.35
72.26
70.86
70.34
69.64
69.24
67.72
67.09
62.38
38.27
37.62
37.13
37.02
36.29
25.48
25.07
24.46
24.09
21.19
17.84
17.82
17.80
17.01
16.94
-0.01
f1 (ppm)

Landomycin B

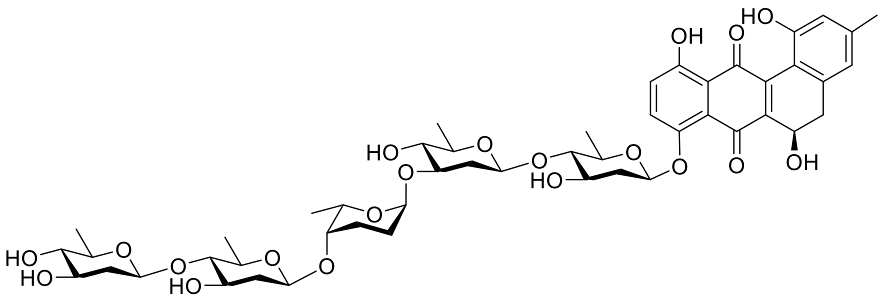

Chemical Formula: $C_{49}H_{64}O_{20}$

$^{1}$H NMR (500 MHz, DMSO) δ 11.69 (s, 1H), 9.63 (s, 1H), 7.51 (d, *J* = 9.4 Hz, 1H), 7.32 (d, *J* = 9.4 Hz, 1H), 6.63 (s, 1H), 6.57 (s, 1H), 5.25 (d, *J* = 9.4 Hz, 1H), 5.11 (d, *J* = 5.6 Hz, 1H), 5.04 (m, 2H), 4.96 (m, 2H), 4.88 (s, 1H), 4.77 – 4.63 (m, 2H), 4.59 (m, 2H), 4.52 (d, *J* = 9.6 Hz, 1H), 4.13 (q, *J* = 7.0 Hz, 1H), 3.66 – 3.41 (m, 5H), 3.37 (m, 2H), 3.26 (dt, *J* = 9.1, 6.2 Hz, 2H), 3.10 (t, *J* = 8.8 Hz, 1H), 2.94 (m, 2H), 2.86 (d, *J* = 16.1 Hz, 1H), 2.81 – 2.68 (m, 2H), 2.37 (dd, *J* = 38.5, 9.6 Hz, 2H), 2.25 (s, 3H), 2.15 – 2.00 (m, 2H), 1.93 (m, 1H), 1.88 – 1.72 (m, 2H), 1.66 (q, *J* = 11.3 Hz, 1H), 1.40 – 1.26 (m, 4H), 1.19 (dt, *J* = 21.7, 5.6 Hz, 12H), 1.00 (d, *J* = 6.4 Hz, 3H).

$^{13}$C NMR (126 MHz, DMSO) δ 188.33, 181.48, 155.80, 155.45, 149.76, 142.22, 141.60, 140.77, 138.74, 128.73, 125.14, 121.45, 119.74, 116.34, 115.91, 113.66, 101.24, 100.67, 100.39, 98.17, 92.55, 87.57, 86.90, 76.86, 76.11, 74.33, 73.60, 72.39, 70.62, 70.51, 70.14, 69.14, 68.78, 65.66, 57.81, 39.98, 39.21, 38.65, 37.08, 36.22, 24.47, 21.55, 18.36, 18.23, 18.19, 18.12, 17.41.

ESI-MS: [M+Na]: 995.36

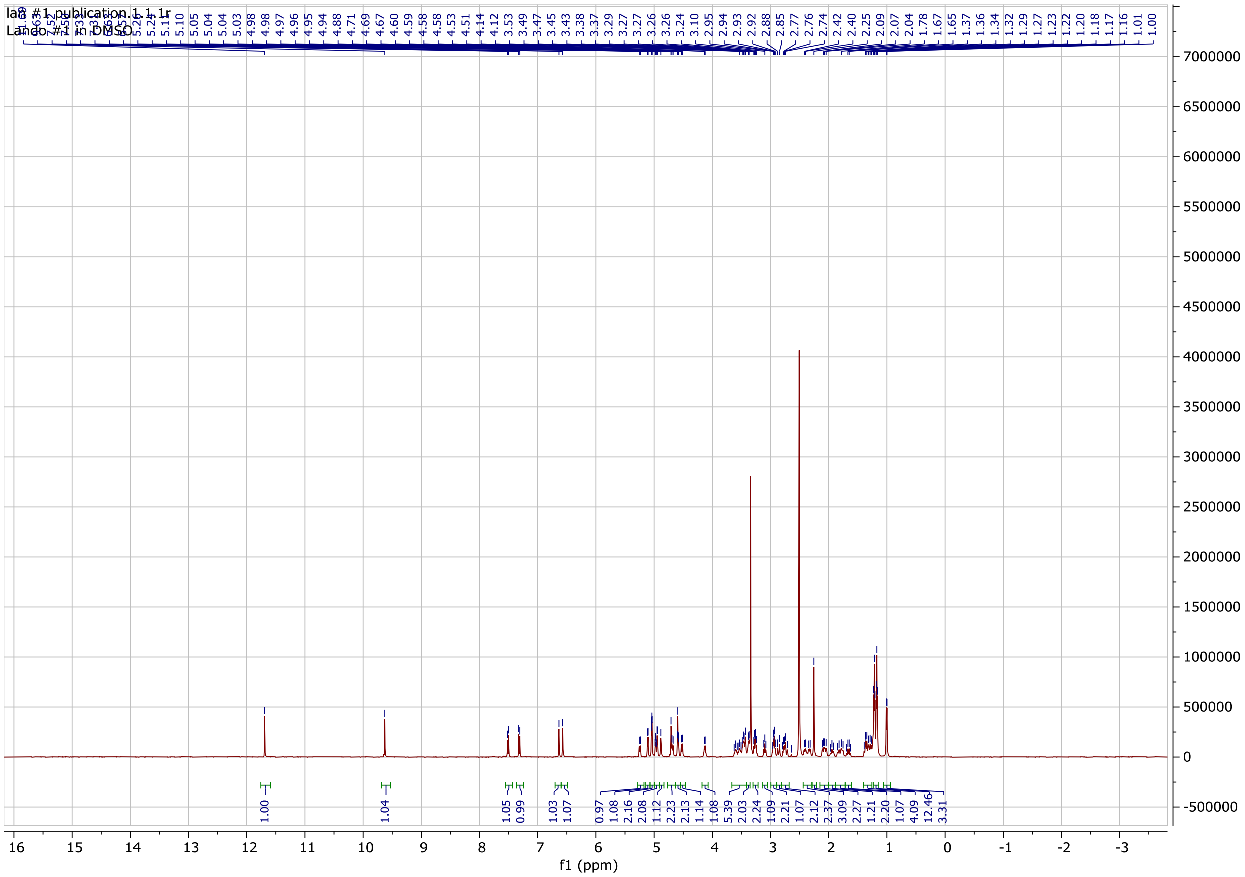
lab #1 publication.1.1.1r
Lando #1 in DMSO
f1 (ppm)

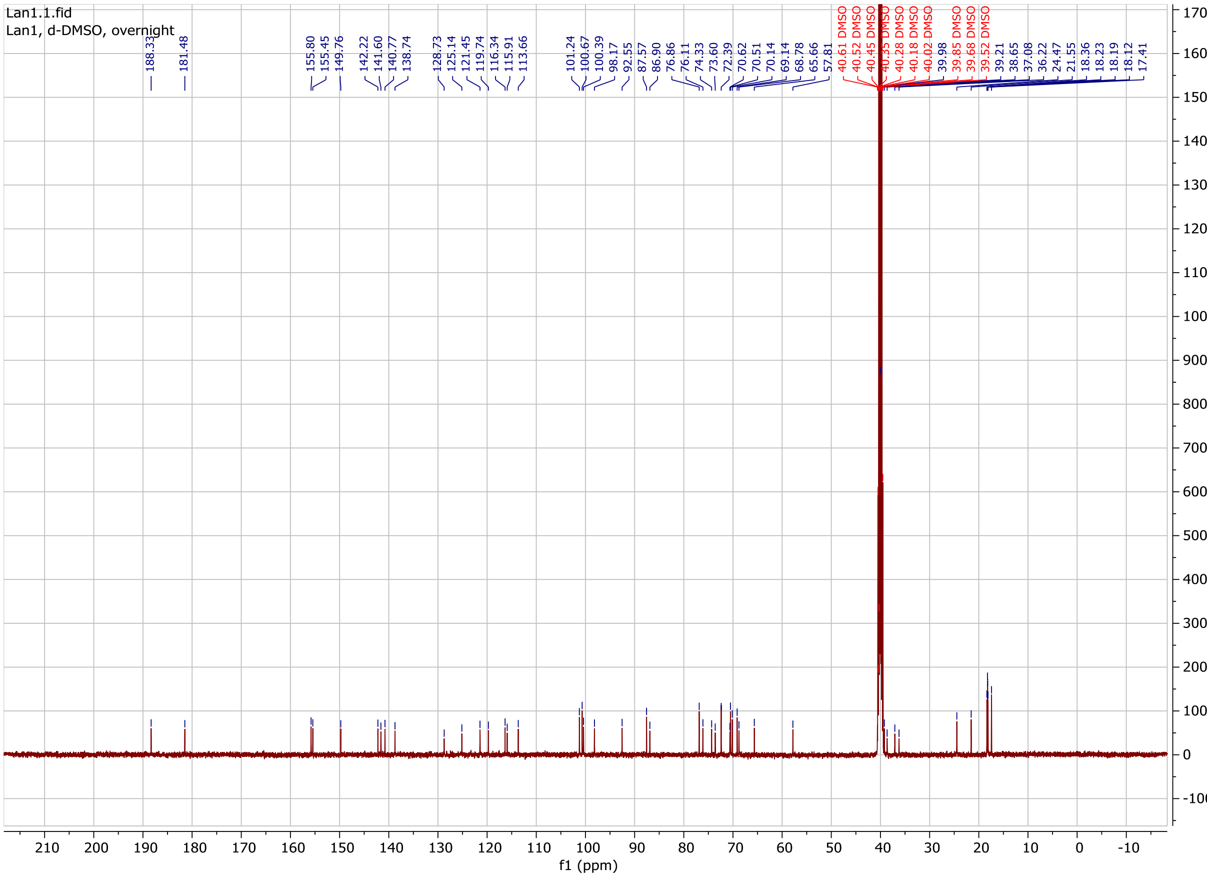
Lan1.1.fid
Lan1, d-DMSO, overnight
188.33
181.48
155.80
155.45
149.76
142.22
141.60
140.77
138.74
128.73
125.14
121.45
119.74
116.34
115.91
113.66
101.24
100.67
100.39
98.17
92.55
87.57
86.90
76.86
76.11
74.33
73.60
72.39
70.62
70.51
70.14
69.14
68.78
65.66
57.81
40.61 DMSO
40.52 DMSO
40.45 DMSO
40.35 DMSO
40.28 DMSO
40.18 DMSO
40.02 DMSO
39.98
39.85 DMSO
39.68 DMSO
39.52 DMSO
39.21
38.65
37.08
36.22
24.47
21.55
18.36
18.23
18.19
18.12
17.41
f1 (ppm)

Landomycin D

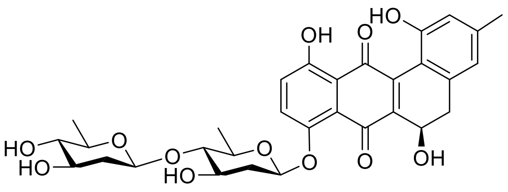


Chemical Formula: $C_{31}H_{34}O_{12}$

[1]H NMR (500 MHz, DMSO) δ 11.69 (s, 1H), 9.63 (s, 1H), 7.51 (d, *J* = 9.4 Hz, 1H), 7.32 (d, *J* = 9.3 Hz, 1H), 6.64 (s, 1H), 6.57 (s, 1H), 5.35 – 5.17 (m, 1H), 5.10 – 4.88 (m, 4H), 4.71 (s, 1H), 4.63 (d, *J* = 9.6, 1H), 3.59 (m, 1H), 3.47 (dd, *J* = 9.1, 6.0 Hz, 1H), 3.25 (s, 1H), 3.07 (t, *J* = 8.9 Hz, 1H), 2.92 – 2.68 (m, 3H), 2.44 – 2.35 (m, 1H), 2.26 (s, 3H), 2.07 (dd, *J* = 13.3, 5.1 Hz, 1H), 1.66 (d, *J* = 11.5 Hz, 2H), 1.36 (m, 1H), 1.20 (d, *J* = 6.2 Hz, 6H)

ESI-MS: [M+Na]: 621.27

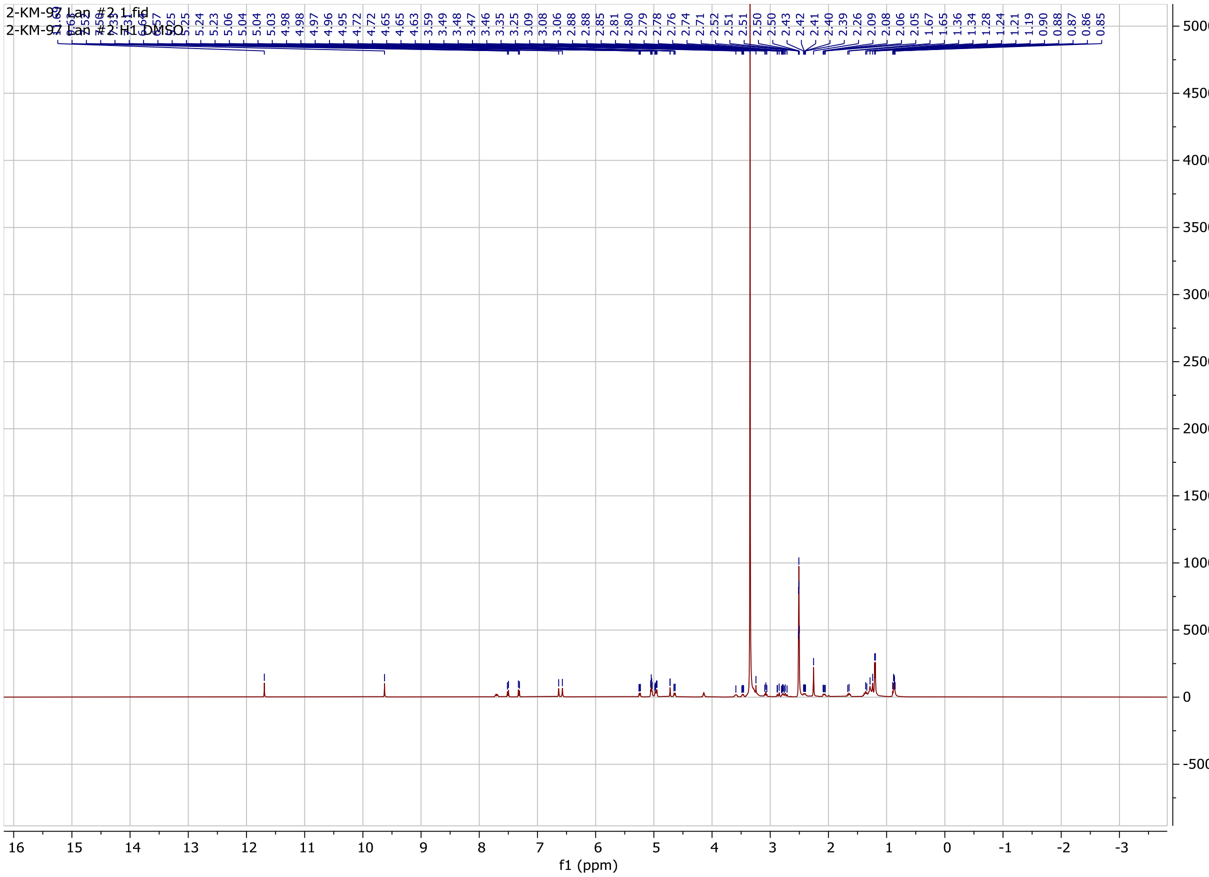

Landomycinone

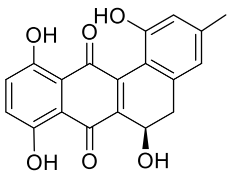

Chemical Formula: $C_{19}H_{14}O_6$

$^{1}$H NMR (500 MHz, $CDCl_3$) δ 12.90 (s, 1H), 12.80 (s, 1H), 9.04 (s, 1H), 7.42 – 7.31 (m, 2H), 6.85 (s, 1H), 6.80 (s, 1H), 5.26 (m, 1H), 3.14 (dd, *J* = 15.8, 4.8 Hz, 1H), 2.97 (dd, *J* = 15.6, 4.3 Hz, 1H), 2.67 (d, *J* = 5.4 Hz, 1H), 2.36 (s, 3H).

ESI-MS: [M+Na]: 360.19

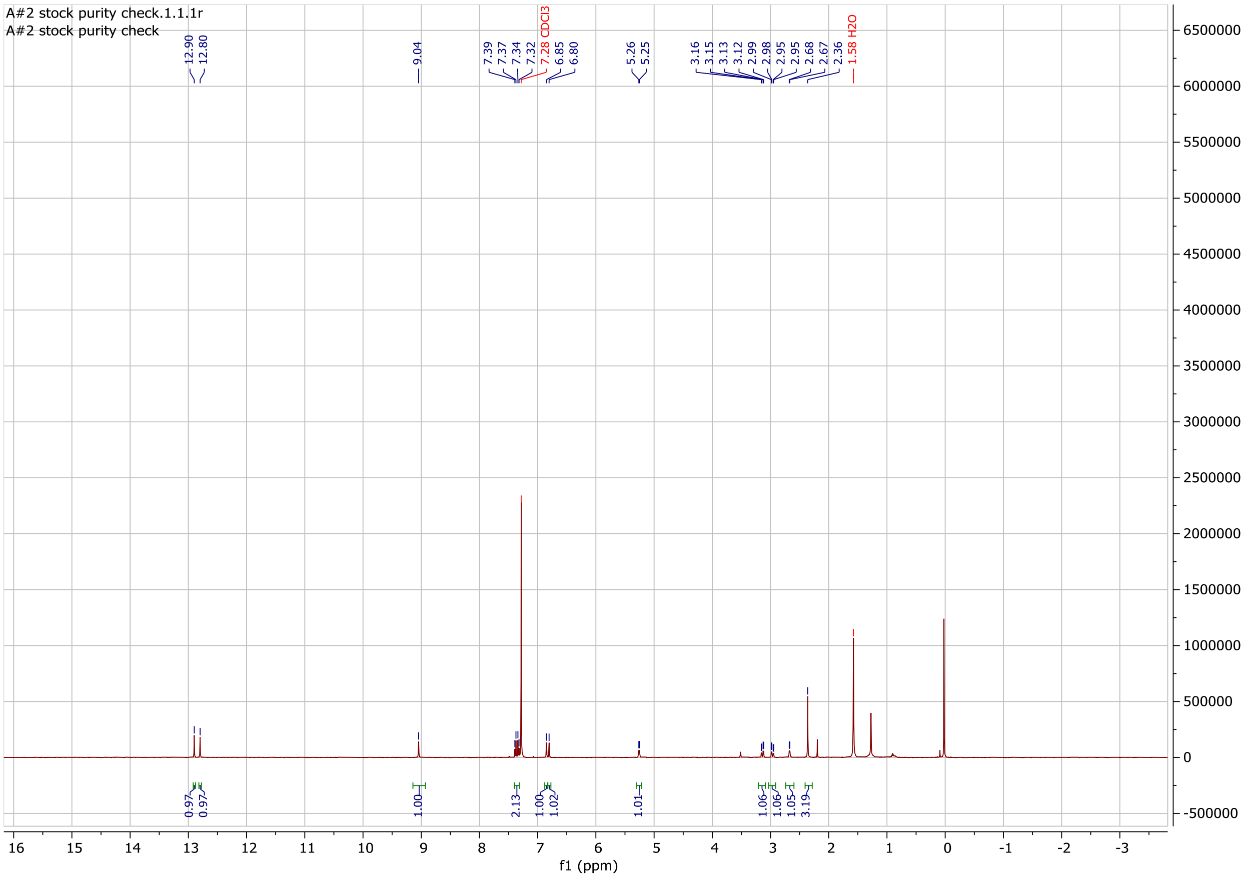

Anhydrolandomycinone

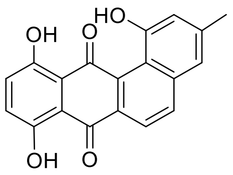

Chemical Formula: $C_{19}H_{12}O_5$

[1]H NMR (500 MHz, $CDCl_3$) δ 13.04 (s, 1H), 12.55 (s, 1H), 11.15 (s, 1H), 8.39 (d, *J* = 8.6 Hz, 1H), 8.20 (d, *J* = 8.6 Hz, 1H), 7.44 – 7.34 (m, 2H), 7.33 (s, 1H), 7.22 (d, *J* = 1.9 Hz, 1H), 2.54 (s, 3H).

ESI-MS: [M+Na]: 342.01

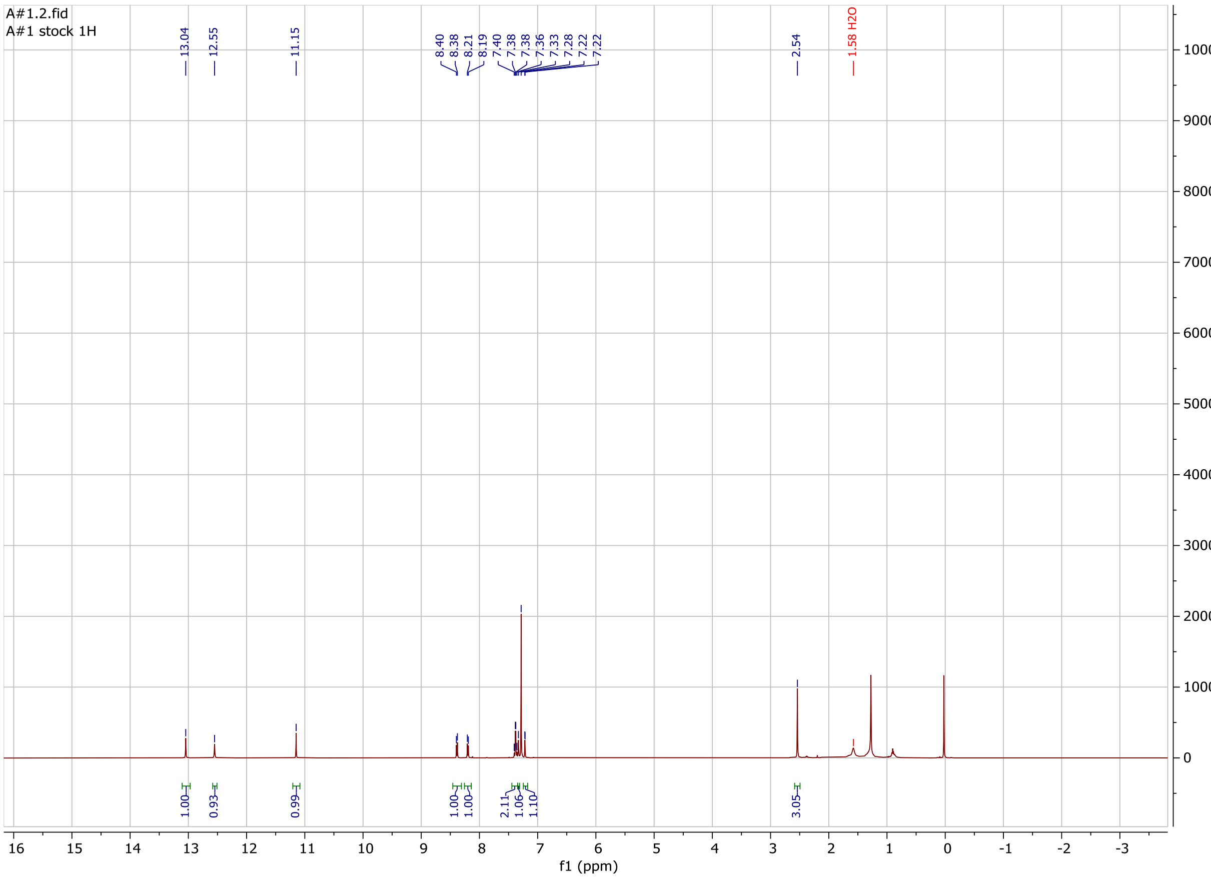

Compound structures were verified with previously reported data.[2–6]

#### References


1. Lambert, R. J. W. & Pearson, J. Susceptibility testing: accurate and reproducible minimum inhibitory concentration (MIC) and non-inhibitory concentration (NIC) values. *J. Appl. Microbiol.* **88**, 784–790 (2000).

2. Yang, X., Fu, B. & Yu, B. Total synthesis of landomycin A, a potent antitumor angucycline antibiotic. *J. Am. Chem. Soc.* **133**, 12433–12435 (2011).

3. Henkel, T., Rohr, J., Beale, J. M. & Schwenen, L. Landomycins, new angucycline antibiotics from streptomyces sp. I. Structural studies on landomycins A~D. *J. Antibiot. (Tokyo).* **43**, 492–503 (1990).

4. Weber, S., Zolke, C., Rohr, J. & Beale, J. M. Investigations of the biosynthesis and structural revision of landomycin A. *J. Org. Chem.* **59**, 4211–4214 (1994).

5. Yang, X. & Yu, B. Synthesis of Landomycin D: Studies on the Saccharide Assembly. *Synthesis (Stuttg).* **48**, 1693–1699 (2016).

6. Roush, W. R. & Neitz, R. J. Studies on the Synthesis of Landomycin A. Synthesis of the Originally Assigned Structure of the Aglycone, Landomycinone, and Revision of Structure. *J. Org. Chem.* **69**, 4906–4912 (2004).
